# A GPNMB-associated lysosomal program tracks neurodegeneration across the DAM continuum

**DOI:** 10.64898/2026.11.20.767890

**Authors:** Georg Werner, Jonas Schulte-Schrepping, Ramona Rodde, Johannes Gnörich, Elena De Domenico, Georg Kislinger, Karsten Nalbach, Lina Dinkel, Johannes C.M. Schlachetzki, Selina Hummel, Astrid F. Feiten, Patricia Hoffelner, Brigitte Nuscher, Marvin Reich, Iñaki Paris, Samira Parhizkar, Alessio Colombo, Stephan A. Müller, Michael Willem, Sven Lammich, Christian Schrader, Jannis B. Spintge, Danilo Prtvar, Alessio Ricci, Stefan Roth, Matthew J. Simon, Todd Logan, Benedikt Wefers, Wolfgang Wurst, Regina Feederle, Arthur Liesz, Dieter Edbauer, Marc D. Beyer, Antje Grosche, Kohichi Kawahara, Christopher Glass, Gilbert Di Paolo, Sabina Tahirovic, Stefan F. Lichtenthaler, Mikael Simons, Martina Schifferer, Matthias Brendel, Christian Haass, Anja Capell, Joachim L. Schultze

**Author notes:** shared corresponding authors Anja Capell Joachim L. Schultze. equal contribution (Georg Werner, Jonas Schulte-Schrepping).

## Abstract

Frontotemporal dementia (FTD) and Alzheimer’s disease (AD) profoundly remodel microglial states, yet how the heterogeneity of disease-associated microglia (DAM) relates to neurodegeneration remains incompletely understood. Using a progranulin (PGRN) and TMEM106B-deficient mouse model recapitulating key features of frontotemporal lobar degeneration (FTLD), including lysosomal impairment, we demonstrated that CSF1R inhibition reduced microgliosis yet accelerated neurodegeneration, suggesting a protective microglia contribution under lysosomal stress. Single-cell transcriptomics revealed that increasing *Gpnmb* expression within DAM was associated with a shift from immune and inflammatory processes toward a lysosomal, autolysosomal, and lipid-catabolic program. Across disease models, engagement of the GPNMB-associated program correlated with neurodegeneration. Consistently, Aβ-driven models with limited neurodegeneration exhibited prominent DAM activation but comparatively limited GPNMB expression and engagement of this program. The GPNMB-associated program was TREM2-dependent, while loss of TREM2 signaling under lysosomal stress was associated with exacerbated neuronal damage. Conversely, lysosomal rescue markedly reduced the GPNMB-associated program, demonstrating its reversibility. CSF proteomics identified a convergent lysosomal and microglial signature across models overlapping with alterations reported in human FTD. Together, GPNMB marks reversible, TREM2-regulated lysosomal transcriptional adaptation along the DAM continuum that scales with lysosomal burden and neurodegenerative severity, consistent with a potentially protective microglial response under lysosomal stress.

## Introduction

Microglia are the innate immune cells of the central nervous system (CNS) and serve essential roles in development, tissue surveillance, and maintenance of neuronal function. In neurodegenerative diseases such as Alzheimer’s disease (AD) and frontotemporal dementia (FTD), microglia undergo pronounced transcriptional and functional reprogramming ^1–4^. Across diverse models of CNS disorders and aging, a distinct population of microglia has been observed often localizing to sites of neuronal stress and amyloid β-peptide (Aβ) pathology ^2,5,6^. These disease-associated microglia (DAM) are characterized by downregulation of homeostatic markers (*Tmem119, P2ry12*) and induction of genes linked to lipid metabolism (*Apoe, Lpl*), phagocytosis (*Trem2, Clec7a*), and lysosomal function (*Ctsb, Cd68*) ^2,3,5,6^. Functionally, DAM have been associated with neuroprotective processes, such as synaptic support and debris clearance ^7,8^. However, they have also been linked to neuroinflammatory responses and neuronal injury, and their impact likely depends on disease context and underlying stressor ^1,5^. Consistent with these context-dependent functions, studies employing microglial depletion through CSF1R inhibition have reported divergent outcomes, ranging from reduced Aβ pathology and improved cognitive performance to exacerbated neuronal injury and neurodegeneration ^9–13^. These observations suggest that DAM may not represent a uniform endpoint but rather span a continuum of context-dependent adaptive states shaped by the nature and severity of pathological stress ^14^. In this study, we adopt a framework in which distinct transcriptional programs represent coordinated sets of genes associated with specific cellular processes, whereas microglial states reflect the broader phenotypes arising from their differential engagement. Within this framework, multiple programs may coexist or vary in their relative activity, giving rise to functionally heterogeneous DAM states rather than strictly discrete populations. Defining disease-relevant DAM programs and their associated functional phenotypes is therefore critical for understanding microglial contributions to neurodegeneration and identifying potential therapeutic opportunities 14,15

A regulator of the transition from homeostatic to DAM states is the triggering receptor expressed on myeloid cells (TREM2), a lipid-sensing receptor involved in coordinating metabolic and phagocytic responses to pathological stress ^16^. Loss of TREM2 impairs Aβ clearance and metabolic fitness ^7,17–21^, whereas TREM2 agonism promotes DAM activation and microglial engagement with Aβ-plaques ^22–25^. These findings position TREM2-dependent programs as a key axis of microglial adaptation in neurodegeneration. However, how TREM2-dependent and -independent DAM responses diversify into distinct functional programs, and how these programs relate to the extent of neurodegeneration, remains incompletely understood.

Among the pathological processes that may shape microglial responses, lysosomal dysfunction represents a potentially convergent source of cellular stress across neurodegenerative diseases ^26^. This is particularly apparent in frontotemporal lobar degeneration (FTLD) caused by haploinsufficiency of *GRN*, the gene encoding progranulin (PGRN), and neuronal ceroid lipofuscinosis (NCL) caused by biallelic *GRN* insufficiency, both of which are characterized by lysosomal impairment ^27–31^. PGRN, which is predominantly expressed by microglia, has been implicated in neuroinflammation, wound healing, and lysosomal biogenesis ^32,33^.

Furthermore, genetic variants in TMEM106B, which encodes a lysosomal membrane protein, modify FTLD risk in individuals carrying GRN mutations ^34–37^.

The functional interaction between PGRN and TMEM106B is evident in *Grn^-^*^/-^ x *Tmem106b*^-/-^ (DKO) mice, in which the combined deficiency exacerbates lysosomal dysfunction, impairs degradative capacity, and accelerates neurodegeneration compared with relatively milder phenotypes in *Grn^-/-^* mice ^38–40^. This model therefore provides a disease context in which pronounced lysosomal stress and progressive neurodegeneration coincide, enabling investigation of microglial responses under increasing pathological burden.

GPNMB (glycoprotein non-metastatic melanoma protein B, also known as osteoactivin) has been described as a marker of DAM in Aβ mouse models and sporadic cases of Alzheimer’s disease ^41–43^ More recent studies have now linked GPNMB to lysosomal dysfunction and DAM remodeling. In PGRN-deficient mice, lysosomal stress induced a GPNMB⁺ microglia state regulated through MITF/TFE, with GPNMB contributing to lysosomal acidification, while complementary work identified conserved transcriptional and epigenetic responses in amyloidosis, prions, and mucopolysaccharidosis type IIIA models ^41,44–48^. Consistent with these findings, a human single-cell atlas identified a GPNMB-high disease associated subtype that expands with AD pathology and is maintained by MITF ^49^.

Although DAM activation is observed across diverse neurodegenerative conditions, how microglial responses adapt to disease-specific contexts and which transcriptional programs are associated with the extent of neurodegeneration remains insufficiently resolved. Here, we hypothesized that increasing GPNMB expression marks a shift within the DAM continuum toward a lysosomal and lipid-catabolic program specific to neurodegeneration. We further asked whether attenuating microgliosis under lysosomal stress exacerbates neuronal injury and whether the GPNMB-associated program can be reversed by restoring lysosomal function.

We combined genetic, transcriptomic, proteomic, imaging, and functional approaches across complementary lysosomal and Aβ-driven disease models to investigate how microglial responses to lysosomal stress relate to neurodegeneration in distinct pathological contexts. First, using the *Grn^-^*^/-^ x *Tmem106b*^-/-^ *DKO* mouse model, we identified a GPNMB-associated transcriptional program characterized by increasing engagement of lysosomal and lipid-catabolic processes, further refining microglial adaptation under pronounced lysosomal dysfunction and neurodegeneration. CSF1R inhibition in this context attenuated microgliosis yet accelerated neuronal injury. Comparisons across lysosomal dysfunction models (*Grn^-/-^*, *Npc1^-/-^*, and *Ctsd^-/-^*), Aβ-driven models (APPPS1), and additional disease paradigms ^39,50–53^, revealed that the GPNMB-associated response more closely tracks the extent of neurodegeneration than broad DAM activation. Moreover, the GPNMB-associated program was enriched in regions of pronounced pathology, its induction was regulated by TREM2 signaling, and its engagement was markedly reduced following restoration of lysosomal function. In addition, CSF proteomics revealed a convergent lysosomal signature across disease models that overlapped with alterations reported in human FTD.

## Results

### Attenuating microgliosis by CSF1R inhibition accelerates neurodegeneration in an FTLD-like mouse model

*Grn^-/-^ x Tmem106b^-/-^* (*DKO*) mice exhibit key features of FTLD, including a significantly reduced lifespan, neurodegeneration and deposition of insoluble, hyperphosphorylated TDP-43 ^38–40^. These pathological features are accompanied by pronounced defects in lysosomal and autophagic degradation, as evidenced by the accumulation of ubiquitinated proteins, LC3-II, and p62. Thus, the *DKO* model combines pronounced lysosomal dysfunction with neurodegeneration, providing an opportunity to test how DAM relate to progressive neurodegeneration.

To investigate the contribution of microgliosis to FTLD-like pathology driven by lysosomal stress, we treated *DKO* and wild-type (*WT*) mice with the CSF1R inhibitor PLX5622 to deplete microglia starting at two months of age for eight weeks (Figure 1a). At this early timepoint, *DKO* mice exhibited no prominent motor phenotype ^39,54^. TSPO-PET imaging at 3.5 months of age revealed a significant signal increase in *DKO* animals compared to *WT* mice, consistent with widespread microglial activation throughout the brain, most prominently in the brain stem (fold change of 1.7) and to a lesser extent in the cortex (fold change of 1.4) (Figure 1b, c). CSF1R inhibition substantially reduced TSPO-PET signal in *WT* and *DKO* mice, although the signal remained above *WT* level in PLX5622-treated *DKO* mice, suggesting residual microglia activity (Figure 1b, c).

**Fig. 1:**
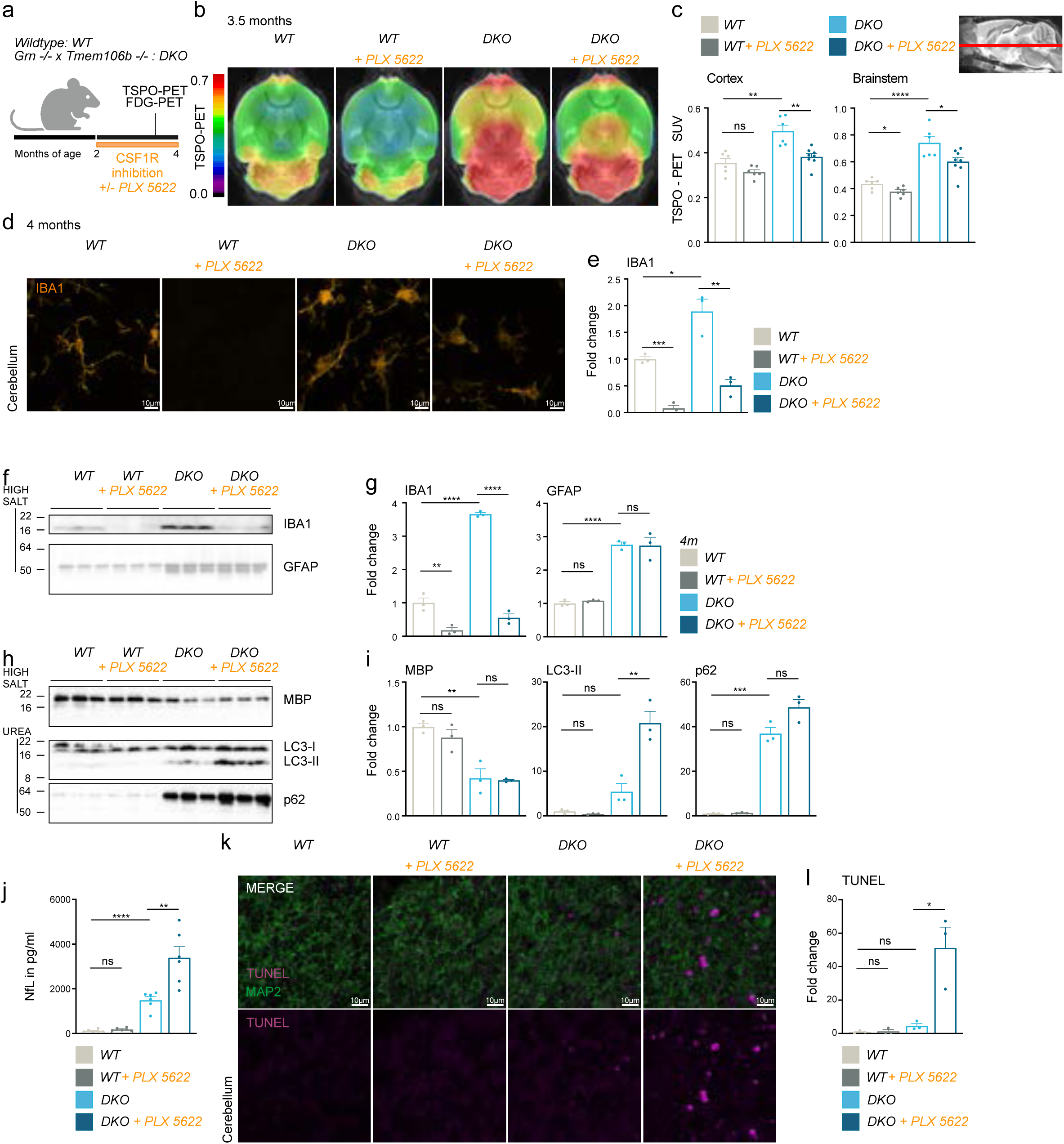
Attenuating microgliosis by CSF1R-inhibition accelerates neurodegeneration in an FTLD-like mouse model. (a) Study overview, CSF1R-inhibition by PLX5622 on *wildtype* (*WT*) mice and *Grn^-/-^* /*Tmem106b^-/-^*(*DKO*) mice starting at 2 months of age for a duration of 2 months. (b) *In vivo* ^18^F-GE180 TSPO µPET imaging. *WT* and *DKO* mice underwent TSPO µPET at 3.5-months of age. All analyses were performed by PMOD (V3.5, PMOD technologies). Normalization of injected activity was performed by the previously validated myocardium correction method ^112^. (c) Quantification of *in vivo* ^18^F-GE180 TSPO µPET imaging. Data represent the mean Standardized Uptake Value (SUV) ± SD (n = 5 - 6 mice per group). (d) Immunofluorescence of IBA1 in the cerebellum of sagittal brain sections. Representative images of 4-month-old *WT* and *DKO* mice ± PLX5622 treatment. Scale bar indicates 10 µm. (e) Quantification of IBA1-immunofluorescence, displayed as fold change to *WT* (n = 3 mice per group).(f) Western blot analysis of IBA1 and GFAP in HIGH SALT (HS) fraction of total brain lysates from of 4-month-old *WT* and *DKO* mice ± PLX5622 treatment, (n = 3 mice per group). (g) Quantification of proteins in (F), fold change to *WT* (n = 3 mice per group). (h) Western blot analysis of MBP in HS-fraction and LC3 and p62 in RIPA-insoluble UREA-fraction of total brain lysates, from of 4-month-old *WT* and *DKO* mice ± PLX5622 treatment, (n = 3 mice per group). (i) Quantification of proteins in (H), fold change to *WT* (n = 3 mice per group). (j) NfL in pg/ml of plasma of mice at 4 months (n = 5 - 6 mice per group). (k) Analysis of apoptotic cells by TUNEL-immunofluorescence in the cerebellum, displayed as magenta. Representative images of 4-month-old *WT* and *DKO* mice ± PLX5622 treatment. Scale bar indicates 10 µm. MAP2 in green as control. (l) Quantification of TUNEL-immunofluorescence, displayed as fold change to *WT* (n = 3 mice per group). (m)*I* Data information: data are depicted as mean ± SD, data points represent individual animals, one-way Anova with Tukey’s multiple comparison was performed. ns, *P < 0.05; **P < 0.01; ***P < 0.001; and ****P < 0.0001.

We next assessed microglial depletion histologically and biochemically. IBA1 immunofluorescence analysis at 4 months of age confirmed successful microglia depletion in *WT* mice treated with PLX5622 and a marked attenuation of microgliosis in *DKO* mice (Figure 1d, e). Consistently, immunoblot analysis of total brain lysates revealed a strong induction of IBA1 in *DKO* mice that was reduced by CSF1R inhibition (Figure 1f, g). GFAP levels, reflecting astrogliosis, were not significantly altered by CSF1R inhibition (Figure 1f, g). Similarly, the reduction in myelin basic protein (MBP) observed in *DKO* animals was not improved (Figure 1h, i). Strikingly, PLX5622-treated *DKO* mice showed an increased accumulation of p62 and LC3-II in the RIPA insoluble fraction (Figure 1h, i). Therefore, CSF1R-inhibition in *DKO* mice is associated with a further increase in autophagy-associated proteins, consistent with aggravated stress.

Next, we investigated whether microglia depletion alters neuronal injury by measuring plasma neurofilament light chain (NfL), a biomarker of axonal damage ^55,56^. NfL levels were significantly elevated in *DKO* mice and further increased upon PLX5622 treatment, with no such effect observed in *WT* mice (Figure 1j). Furthermore, TUNEL staining revealed apoptotic cells in the cerebellum of *DKO* mice, which was markedly exacerbated in PLX5622-treated *DKO* mice (Figure 1k, l). To assess brain glucose uptake and cellular metabolism *in vivo,* we performed FDG-PET. Consistent with prior observations ^57^, microglia depletion in *WT* mice reduced FDG uptake (Figure 1m, n), highlighting the combined metabolic contribution of neurons and microglia. Importantly, PLX5622-treated *DKO* mice exhibited a pronounced reduction in FDG uptake in the brainstem, even though microglia depletion in this region was less complete, and TSPO-PET signal remained elevated (Figure 1c, m, n). Thus, in *DKO* mice, the strongly reduced FDG-PET signal might indicate reduced neuronal metabolic activity or neuronal loss in the brainstem. Together with the increased NfL and TUNEL signals, brainstem hypometabolism further indicated aggravated pathology following CSF1R inhibition.

Collectively, these data show that, while CSF1R inhibition reduces microgliosis, depletion of PLX5622-sensitive microglia worsens neurodegeneration in *DKO* mice, suggesting that microglia may play a protective function in neurodegenerative conditions under lysosomal stress.

### Neurodegeneration-associated microglial subpopulations expand in *DKO* mice and show differential sensitivity to CSF1R inhibition

To define cell type and state-specific responses to genetically imposed lysosomal burden and progressive neurodegeneration, we performed microwell-based single-cell RNA sequencing (scRNA-seq) using the BD Rhapsody platform. Microglia (CD11B⁺), astrocytes (ACSA-2⁺), and oligodendrocytes (O4⁺) were isolated by Magnetic Cell Separation (MACS) and fluorescence-activated cell sorting (FACS) from *WT* and *DKO* mice, with or without CSF1R inhibition via PLX5622 treatment.

After quality control and filtering, we obtained 48,151 high-quality single-cell transcriptomes. These were clustered into 17 transcriptionally distinct populations using Louvain clustering and visualized in two-dimensional space using Uniform Manifold Approximation and Projection (UMAP) (Figure 2a, b).

**Fig. 2:**
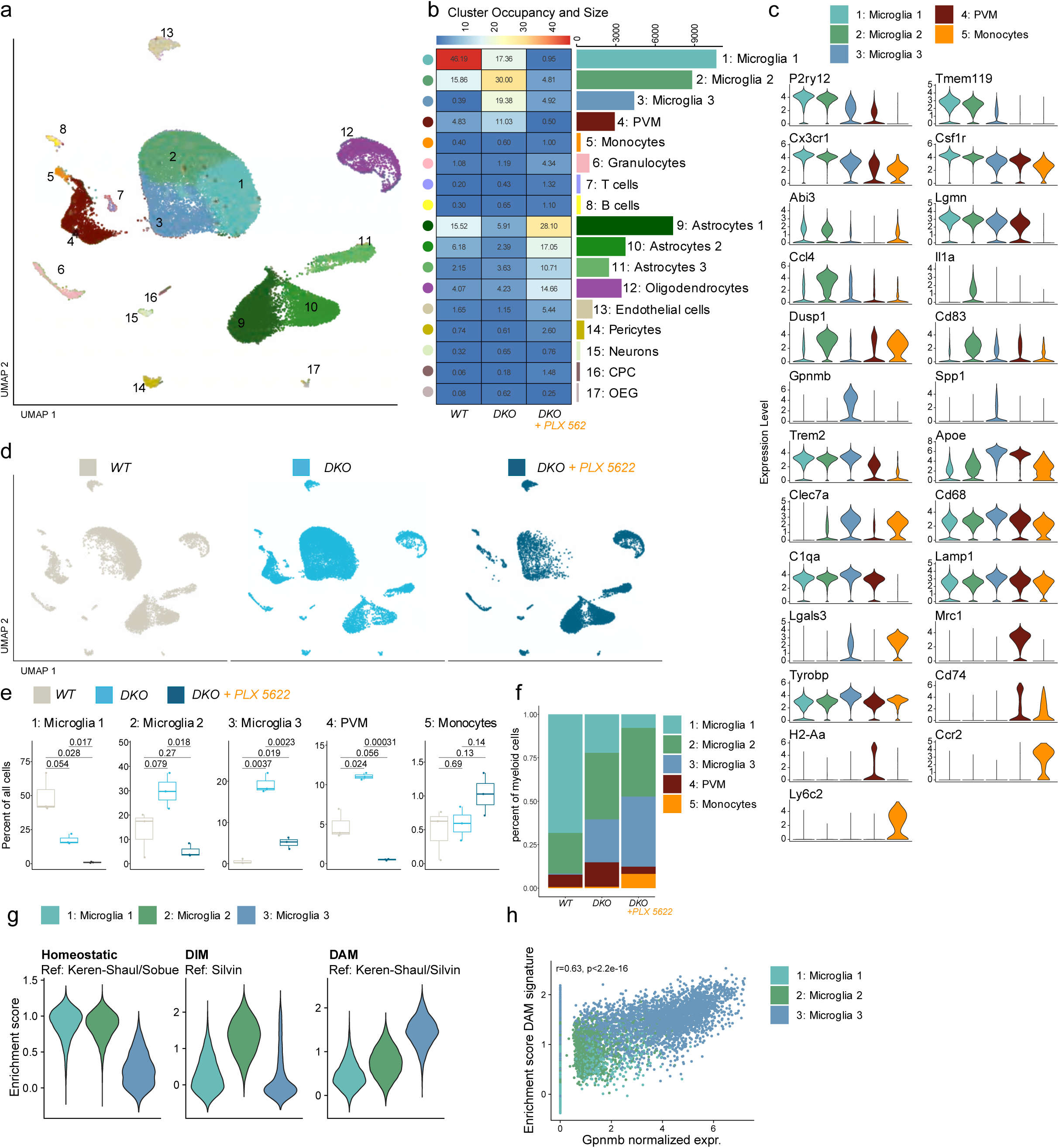
Neurodegeneration-associated microglial subpopulations expand in *DKO* mice and show differential sensitivity to CSF1R inhibtion. (a) UMAP visualization of scRNA-seq profiles of 48,151 cells from *WT* and *DKO* mice (n=3) treated with PLX5622. Cells are clustered and colored according to assigned cell type identities. (b) Heatmap illustrating the proportional contributions of individual conditions across cell types (left). Bar chart showing total number of cells per cluster (right). (c) Violin plot showing expression values of selected marker genes across clusters. (d) UMAP visualizations split and colored according to genotype and treatment. (e) Box plots showing cluster occupancy split by genotype and treatment, as percentage of all cells. P-values of two-sided Student’s *t*-test are displayed. (f) Stacked bar charts of cluster percentages within myeloid cells per genotype and treatment. (g) Violin plots of enrichment values per cell for HOM, DIM and DAM microglia signatures derived from ^2,3,88,113^ (h) Scatter plot of GPNMB expression per cell and DAM enrichment scores of microglia clusters 1, 2 and 3.

Cell type identities were assigned based on canonical marker gene expression (Figure 2c, Figure S1a) and refined by label transfer from published mouse brain single-cell atlases ^58,59^. The majority of captured cells were identified as microglia, astrocytes, and oligodendrocytes, with minor populations of perivascular macrophages (PVMs), monocytes, granulocytes, T cells, B cells, and endothelial cells (Figure 2b).

ScRNA-seq revealed a striking expansion of distinct microglia populations in *DKO* mice (Figure 2d, e, f). Cluster occupancy analysis indicated significant remodeling of the myeloid compartment: Cluster 2 was strongly expanded, while cluster 3 emerged exclusively in *DKO* animals (Figure 2d, e, f). CSF1R inhibition differentially affected these clusters. Clusters 1 and 2 were highly susceptible to depletion, whereas cluster 3 exhibited partial resistance to PLX5622 treatment and remained elevated compared to *WT* (Figure 2d, e, f).

Transcriptomic comparison with established microglia gene signatures revealed that clusters 1 and 2 were enriched for homeostatic microglia genes (Figure 2g, Figure S1b). Cluster 1, characterized by highest expression of *P2ry12* and *Tmem119* (Figure 2c), was designated as homeostatic microglia (HOM) ^2^. Cluster 2 marker expression overlapped significantly with the disease-inflammatory monocyte/macrophage (DIM) signature (Figure 2g), and expressed pro-inflammatory genes such as *Cd83*, *Ccl4*, *Dusp1* and *Il1a* (Figure 2c) ^3^. Disease-associated microglia (DAM) markers, including *Trem2* and *Clec7a* ^2,3^, were differentially expressed across clusters, with the highest expression in cluster 3 (Figure 2c).

Violin plots further demonstrated selective upregulation of *Gpnmb* and *Spp1* (Secreted Phosphoprotein 1, osteopontin) in cluster 3, indicating a disease-associated microglial state characterized by increased lysosomal engagement and phagocytic activity (Figure 2c). *Gpnmb* is frequently elevated in conditions of inflammation and elevated under lysosomal and degradative demand ^47,48,60,61^. *Spp1* has been shown to promote microglial phagocytic states and synaptic engulfment in the context of Aβ pathology linking this DAM cluster to active tissue remodeling and clearance processes (Figure S1C) ^61,62^. Consistent with this interpretation, enrichment analysis showed that cluster 3 closely aligned with DAM gene signatures, and *Gpnmb* expression strongly correlated with overall DAM enrichment scores (Figure 2g, h, S1b). Notably, GPNMB⁺ DAM exhibited partial resistance to CSF1R inhibition, consistent with a changed trophic dependency on CSF1R signaling ^63–65^, possibly affecting proliferation and survival of microglia, which could further affect the response to injury.

Finally, pathway analysis of cluster-specific gene signatures highlighted distinct molecular functions associated with each microglia cluster (Figure S1c). HOM cluster 1 was enriched for cellular maintenance genes involved in RNA processing, DNA repair, and lipid metabolism. DIM cluster 2, consistent with its expression of inflammatory markers, showed an overrepresentation of genes related to immune responses, activation of myeloid and lymphoid cells, autophagy, and the cellular response to unfolded proteins. In contrast, DAM cluster 3 was defined by gene signatures linked to lysosomal function, detoxification processes, and mitochondrial respiration rather than broad inflammatory signaling (Figure S1c). In the setting of genetically induced lysosomal impairment in *DKO* mice, this transcriptional profile is consistent with a microglial response to increased degradative and metabolic demand.

To further characterize the regulatory basis of this phenotype, transcription factor binding motif analysis of cluster-specific genes predicted MITF/TFE family members as regulators of the DAM-associated lysosomal program (Figure S2a,b). Further, H3K27ac profiling of PU.1⁺ nuclei identified differential enhancer activity at cluster 3-associated loci, including *Gpnmb* and *Fabp5* (Figure S2c-e), while motif enrichment in differentially acetylated regions implicated MITF/TFE-associated regulation (Figure S2f). These findings support recent reports linking lysosomal dysfunction to MITF/TFE-associated transcriptional and epigenetic remodeling of GPNMB⁺ microglia ^47,48^.

We further validated the GPNMB⁺ DAM phenotype at the protein and tissue level. GPNMB, SPP1 and GALECTIN3 were strongly increased in *DKO* brain and remained largely preserved following CSF1R inhibition, whereas CLEC7A and TREM2 were induced in DKO mice but substantially reduced by PLX5622 treatment (Figure S3a, b). In contrast, CSF1R and ABI3 were strongly reduced following CSF1R inhibition (Figure S3a, b), consistent with the differential sensitivity of the microglial populations observed by scRNA-seq. Immunofluorescence further confirmed the presence of GPNMB⁺/CLEC7A⁺ microglia in the cortex, cerebellum and thalamus of *DKO* mice and their relative persistence following PLX5622 treatment (Figure S3c).

Together, these data identify cluster 3 as a DAM-enriched, GPNMB⁺/SPP1⁺ microglial population with prominent lysosomal and metabolic programs that shows reduced sensitivity to CSF1R inhibition.

### GPNMB⁺ DAM exhibit lysosomal overload

To determine whether the lysosomal transcriptional program of GPNMB⁺ DAM reflects increased lysosomal demand *in vivo*, we examined microglial lysosomal morphology and function in *DKO* mice. Within the scRNA-seq data, Ctsb and Ctsd were expressed across microglial clusters, with the highest expression in cluster 3 (Figure 3a), further supporting increased lysosomal engagement of this population. Ultrastructural analysis of the thalamus revealed pronounced accumulation of lipid-rich and crystalline material within lysosomal structures of *DKO* microglia, frequently in proximity to dystrophic neurites (Figure 3b). These structures are consistent with previously described crystalline lipid deposits under conditions of microglial lipid overload ^66–69^.

**Fig. 3:**
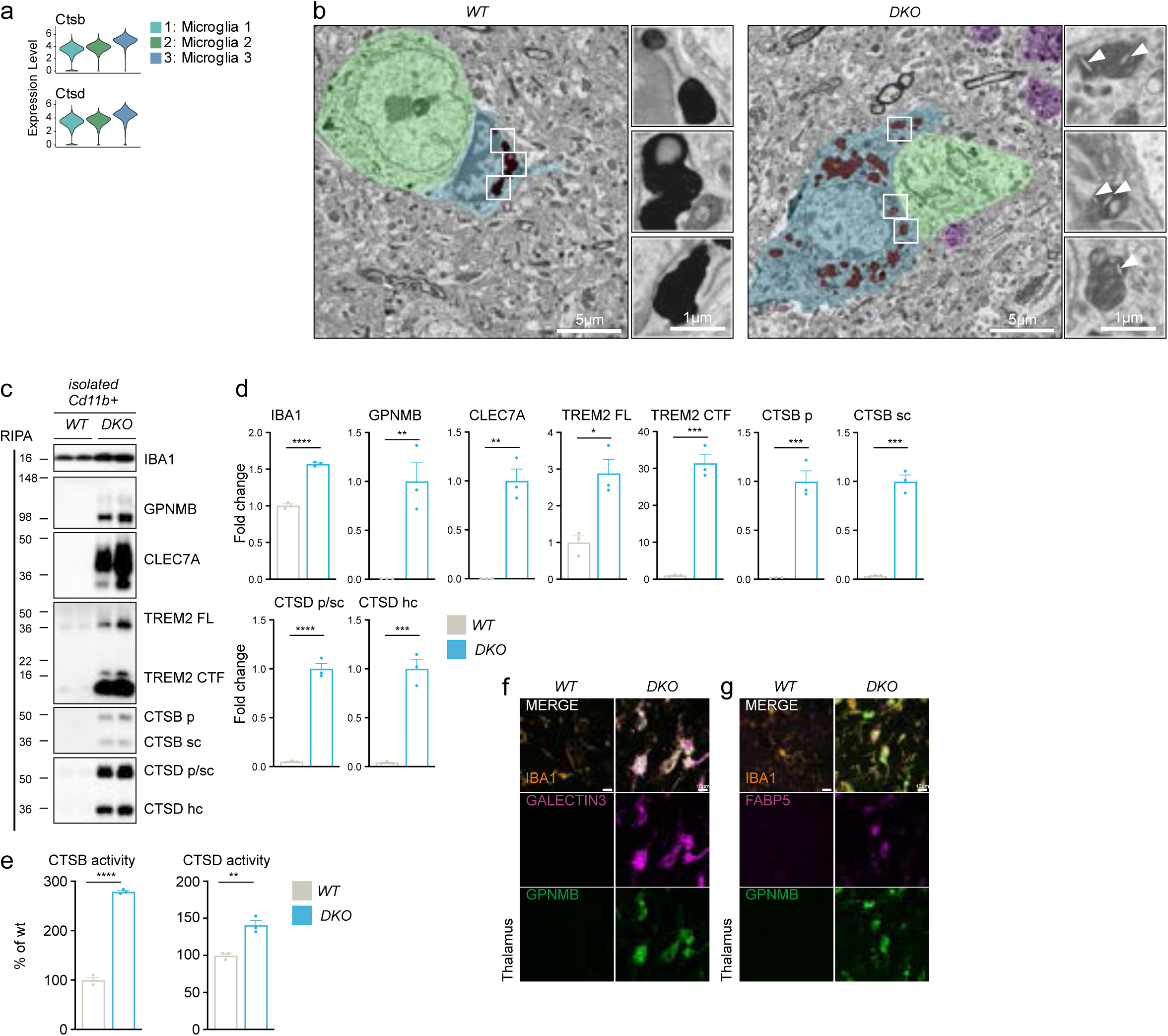
Lysosomal impairment in GPNMB⁺ DAM is associated with crystal accumulation in *DKO* mice. (a) Violin plots showing *Ctsb* and *Ctsd* expression in scRNA-seq data of microglia cluster 1, 2 and 3 as shown in Fig. 2. (b) Scanning electron microscopy (SEM) of microglia in the thalamus, of 2.5-month-old *WT* and *DKO* mice, representative images showing crystal accumulation within lysosomes of microglia in proximity to neuronal debris. Scale bars indicate 5 µm and 1 µm. Structures are highlighted; neuronal cell body (green), dystrophic neurite (purple), microglia (blue), microglial lysosomes (red) and crystals (white arrows). (c) Western blot analysis of isolated CD11B^⁺^ cells for microglia proteins, IBA1, GPNMB, CLEC7A, TREM2, and lysosomal proteins CTSB and CTSD from of 4-month-old *WT* and *DKO* mice, (n = 3 mice per group). Maturation variants are indicated (hc, heavy chain; p, preform; sc, single chain) (d) Quantification of proteins in (B), displayed as fold change to *WT.* Except for GPNMB, CLEC7A, CTSB, and CTSD, protein levels in *WT* mice were of low signal abundance, therefore *DKO* was used as reference. (e) Catalytic activity of cathepsin B and D in total brain lysates from 4-month-old mice were incubated with quenched fluorogenic substrate for CTSB and CTSD. Fluorescence signal was normalized to wild type. Data represent the mean normalized activity ± SD. (f) Immunofluorescence of lysosomal regulator GALECTIN3 (magenta), microglia activation markers GPNMB (green) and IBA1 (orange) in the thalamus of sagittal brain sections. Representative images of 2.5-month-old *WT* and *DKO* mice. Scale bar indicates 10 µm. (g) Immunofluorescence of lipid binding protein FABP5 (magenta), microglia activation markers GPNMB (green) and IBA1 (orange) in the thalamus of sagittal brain sections. Representative images of 2.5-month-old *WT* and *DKO* mice. Scale bar indicates 10 µm.

Analysis of isolated CD11b⁺ microglia further confirmed strong induction of GPNMB, CLEC7A and TREM2 together with increased abundance and maturation of the lysosomal proteases CTSB and CTSD in *DKO* mice (Figure 3c, d). This increase in lysosomal protease abundance occurred alongside increased levels of TREM2 C-terminal fragments (CTF). Increased CTSB and CTSD activity further supported enhanced lysosomal protease activity in the disease context (Figure 3e). Immunofluorescence additionally showed GPNMB⁺ microglia positive for GALECTIN3, a marker associated with lysosomal damage, and co-expression of GPNMB with the lipid metabolism-associated protein FABP5 in the thalamus of *DKO* mice (Figure 3f, g). Together, these findings indicate that the GPNMB-associated lysosomal program is accompanied by increased lysosomal machinery in the face of pronounced lysosomal cargo accumulation, including lipid-rich material, consistent with lysosomal overload in GPNMB⁺ DAM.

### GPNMB expression marks progressive lysosomal adaptation along the DAM continuum

Having established that GPNMB⁺ DAM exhibit substantial lysosomal burden, we next asked how the differential engagement of transcriptional programs shapes DAM states along a gradient of increasing *Gpnmb* expression. Building on our conceptual framework that DAM states reflect the combined activity of distinct transcriptional programs, we investigated whether increasing *Gpnmb* expression is associated with coordinated changes in the functional composition of the DAM response. We therefore stratified microglia cluster 3 according to *Gpnmb* expression, separating *Gpnmb*-negative cells and dividing *Gpnmb*-expressing cells into bins of progressively increasing expression (Figure 4a). To place these changes in the context of the transition from homeostatic to DAM, homeostatic microglia cluster 1 was included as a reference. Genes differentially expressed between homeostatic, GPNMB-low and GPNMB-high microglia were grouped according to their expression pattern, revealing four major transcriptional programs (P1-P4) (Figure 4b).

**Fig. 4:**
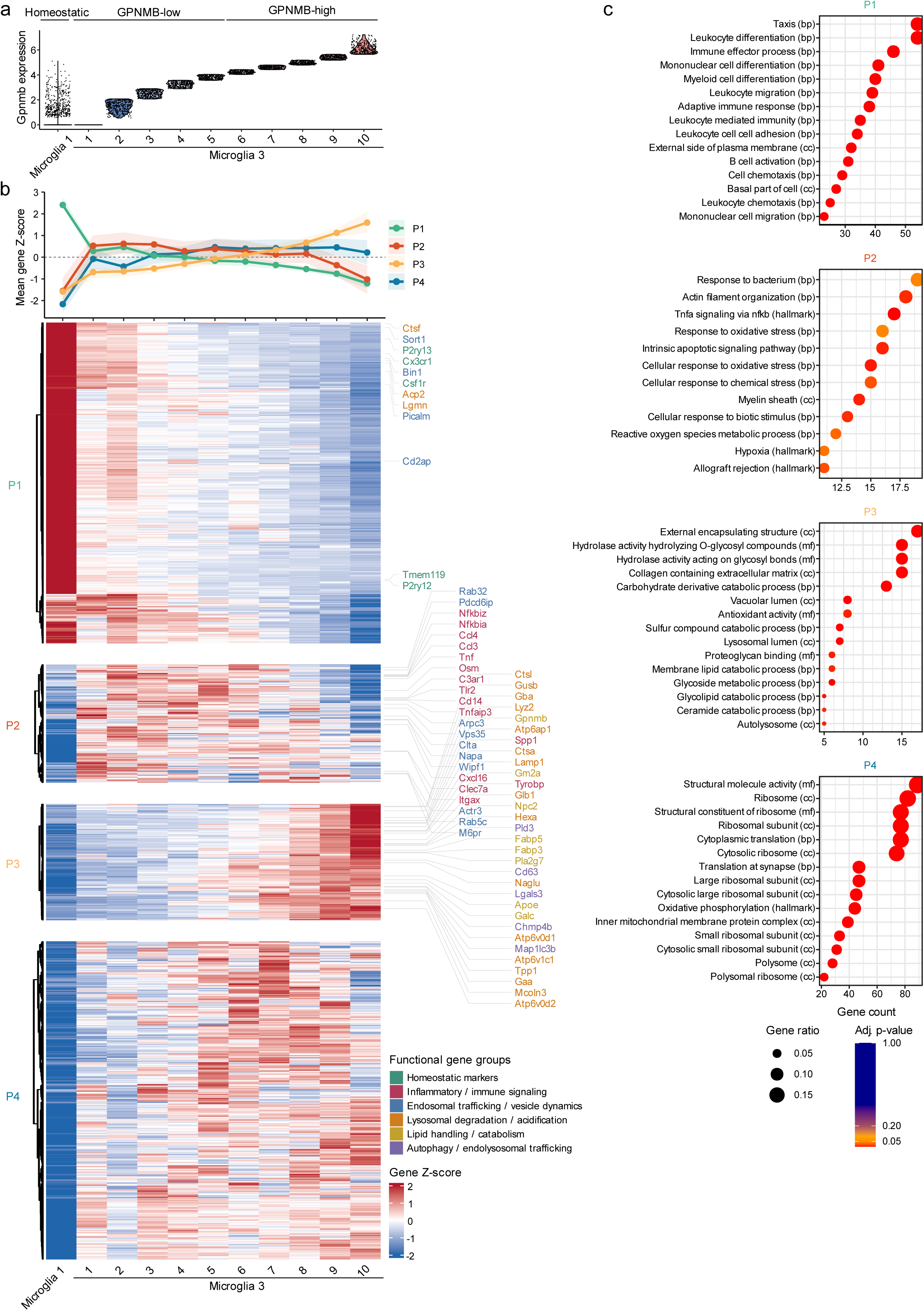
GPNMB⁺ expression marks upregulation of lysosomal processes within the DAM continuum. (a) Violin plot of *Gpnmb* expression across homeostatic microglia (cluster 1) and microglia from cluster 3 binned according to increasing Gpnmb expression. (b) Mean expression trajectories and heatmap of genes differentially expressed between homeostatic, GPNMB-low and GPNMB-high microglia. DAM (microglia 3) were stratified into ten bins of increasing *Gpnmb* expression (see a), and mean gene expression was calculated for each bin. Columns represent homeostatic microglia followed by the ten *Gpnmb* expression bins, and rows represent genes with expression values scaled to gene-wise Z-scores. Hierarchical clustering identified four transcriptional programs (P1–P4) with distinct expression patterns. The line plot above the heatmap shows the mean gene Z-score for each program across the expression gradient, with shaded areas indicating the interquartile range across genes. Selected genes are labeled and color-coded according to their associated biological functions, including homeostatic identity, inflammatory signaling, endosomal trafficking and vesicle dynamics, lysosomal degradation and acidification, lipid handling and catabolism, and autophagy-associated processes. (c) Functional enrichment analysis of genes in P1–P4. Dot size indicates gene ratio, color represents adjusted p-value, and the x-axis indicates the number of genes associated with each enriched term. Enrichment was assessed using Gene Ontology and Hallmark gene sets with Bonferroni correction.

P1 comprised genes most characteristic of homeostatic microglia that were downregulated upon transition to DAM, including genes associated with cell migration and myeloid differentiation, alongside components of endocytic trafficking (*Picalm*, *Bin1*, *Cd2ap*, *Sort1*) (Figure 4b, c). P2 was markedly increased in DAM compared to HOM, but showed a decline with increasing *Gpnmb* expression This program included inflammatory mediators and immune-signaling genes such as *Ccl3*, *Ccl4*, *Tnf* and *Clec7a*, together with components of endosomal trafficking and vesicle dynamics, including *Vps35*, *Rab5c*, *M6pr* and *Pdcd6ip*. In contrast, P3 strongly correlated with increasing *Gpnmb* expression and contained established DAM genes, including *Apoe*, *Lyz2* and *Tyrobp*, alongside a broad repertoire of lysosomal hydrolases (*Ctsl*, *Gba*, *Hexa*, *Gusb*), components of lysosomal acidification and homeostasis (*Atp6ap1*, *Atp6v0d2*, *Lamp1*), and genes involved in lipid handling and catabolism (*Npc2*, *Gm2a*, *Fabp5*) (Figure 4b). Functional enrichment of P3 highlighted lysosomal, autolysosomal and lipid catabolic processes (Figure 4c). P4 was also induced during the transition from homeostatic microglia to DAM but remained largely independent of the *Gpnmb* gradient and was dominated by ribosomal and translation-associated processes.

Together, these findings support a conceptual framework in which DAM states emerge through the differential engagement of transcriptional programs rather than representing strictly discrete populations. Along the *Gpnmb* expression gradient, the relative contribution of these programs shifts from homeostatic and inflammatory responses toward increasing lysosomal and lipid-catabolic activity, while other programs remain largely unchanged. Thus, GPNMB marks a continuum of transcriptional remodeling characterized by progressively increasing engagement of a lysosomal adaptation program within DAM.

### Temporal and regional induction of the GPNMB-associated response tracks neurodegenerative severity

To determine when the GPNMB-associated program emerges during disease progression, we examined *DKO* mice at an early disease stage. At 2 months of age, *DKO* mice already showed increased TSPO-PET signal in cortex and brainstem, indicating early microgliosis, accompanied by elevated plasma NfL levels as a measure of neuronal injury (Figure S4a-c). Analysis of microglial marker genes in the nCounter® Glia Panel data revealed induction of *Gpnmb*, *Spp1*, *Lgals3*, *Clec7a* and *Trem2* at 2 months, with a further pronounced increase by 4 months of age (Figure S4d). Consistent with the transcriptional data, SPP1, GALECTIN3, CLEC7A and TREM2 were increased at the protein level already at this early disease stage (Figure S4e, f). Immunofluorescence further revealed regional accumulation of GPNMB⁺ microglia, particularly in the thalamus (Figure S4g). Together, these findings indicate that the GPNMB-associated program is already induced during early disease and progressively increases alongside neuronal injury.

We next asked whether the association between the GPNMB-associated response and pathological burden was also evident spatially within the brain. We therefore dissected cortex, midbrain and hindbrain at 4 months of age, and assessed microglial activation together with markers of FTLD-like pathology (Figure S5a, c). While increased IBA1 was detected across regions, GPNMB showed a pronounced regional distribution and was most strongly increased in midbrain and hindbrain. These regions also showed the greatest accumulation of p62, phosphorylated TDP-43 and ubiquitinated proteins in the insoluble fraction, whereas cortical changes were comparatively modest (Figure S5c-e). CLEC7A and TREM2 were likewise increased preferentially in the more strongly affected regions.

Together, the temporal and regional analyses show that the GPNMB-associated microglial program emerges early and increases during disease progression, while its regional abundance closely tracks the extent of pathology. These findings further support GPNMB-associated lysosomal remodeling as a feature of DAM that becomes increasingly prominent with advancing neurodegeneration.

### GPNMB-associated lysosomal gene expression correlates with neurodegeneration

To determine whether transcriptional programs identified along the progressive increase in *Gpnmb* expression are associated with the extent of neurodegeneration, we analyzed bulk gene expression in whole brain homogenates using the nCounter® Glia Panel. We included *APPPS1* mice, *Npc1^-/-^* mice, and *DKO* mice at 2 and 4 months of age, representing conditions with different degrees of neuronal injury as reflected by plasma NfL levels, as well as *WT* mice at different ages corresponding to the disease models (Figure 5a). *APPPS1* mice develop pronounced microgliosis with progressive Aβ deposition but comparatively limited overt neurodegeneration, whereas *Npc1^-/-^* mice exhibit defective endolysosomal lipid trafficking and substantial neurodegeneration ^52,53,55,70^. Together with the progressive pathology of *DKO* mice, these models allowed us to compare microglial transcriptional programs across distinct pathological contexts and extent of neurodegeneration. We then assessed enrichment of genes from the P1, P2 and P3 signatures identified in the scRNA-seq analysis that were represented on the nCounter® panel, focusing on the transcriptional programs that showed dynamic changes along the GPNMB-associated trajectory, whereas the largely GPNMB-independent, ribosomal C4 program was not included (Figure 5b).

**Fig. 5:**
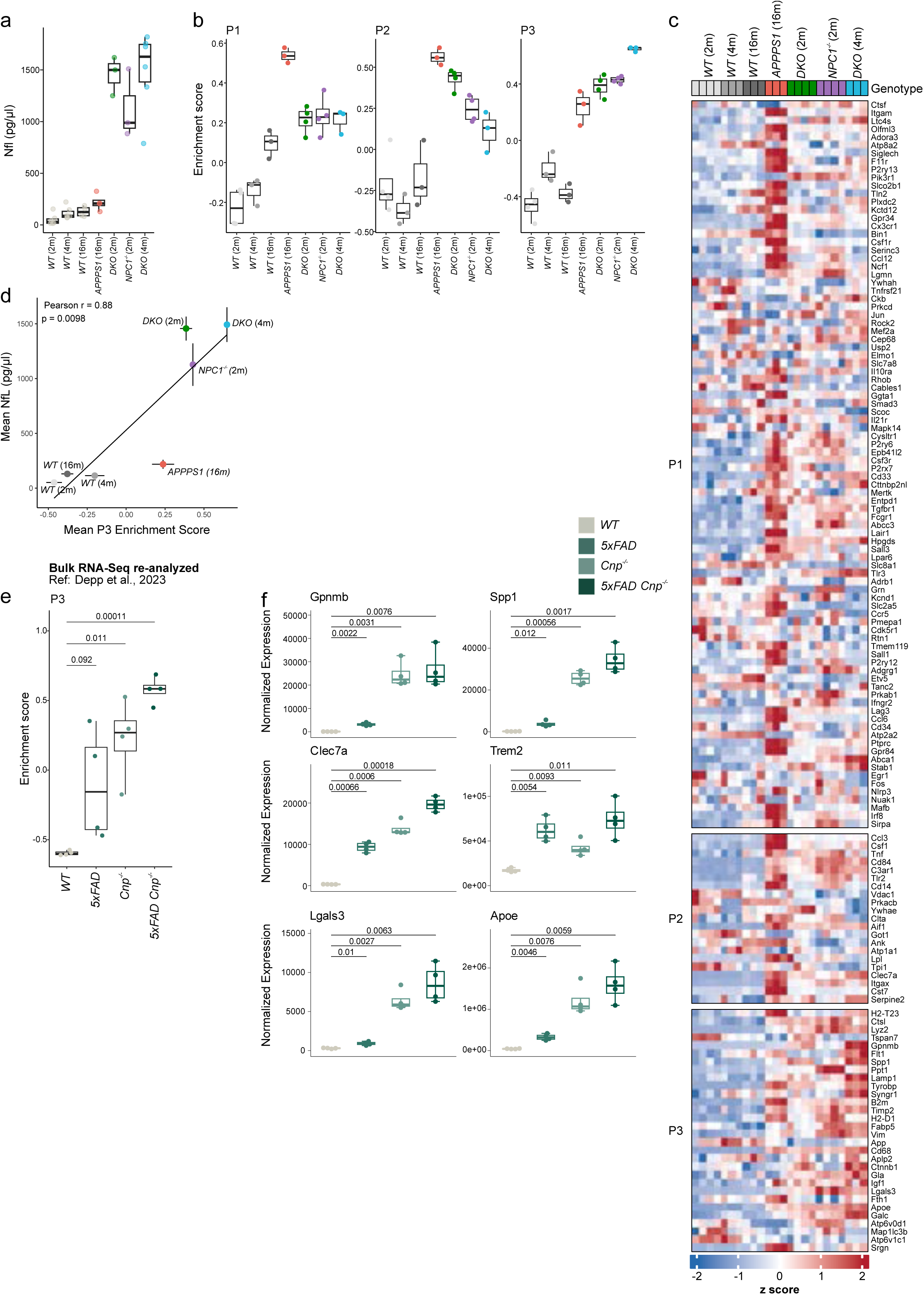
GPNMB⁺ DAM program correlates with neurodegeneration across neurodegeneration mouse models. (a) Box plots showing NfL in pg/ml of plasma of *WT* and *APPPS1*, *DKO* and *Npc1^-/-^* mice at indicated ages, ranging from 2 to 16 months (n = 3 mice per group) (b) Box plots showing enrichment scores for the P1, P2, and P3 microglial programs defined in Figure 4 in bulk brain transcriptomes from *WT, APPPS1*, *DKO*, and *Npc1^-/-^* mice at the indicated ages, ranging from 2 to 16 months. (c) Heatmap showing z-score-scaled expression of P1, P2, and P3 program genes (Fig. 4B) in bulk brain transcriptomes from *WT, APPPS1, DKO*, and *Npc1^-/-^* mice at the indicated ages. (d) Pearson correlation between plasma NfL levels and mean enrichment of the GPNMB-associated P3 signature across the indicated experimental groups. (e) Box plot showing enrichment scores of the P3 program (Fig. 4B) in bulk RNA-seq data from CD11B^⁺^ microglia from 6-month-old *WT, 5xFAD, Cnp^-/-^*and *5xFADxCnp^-/-^* mice ^73^. P-values from two-sided Student’s *t*-tests are shown. (f) Box plots of normalized counts of selected microglia genes in CD11B^⁺^ microglia from *WT, 5xFAD, Cnp^-/-^* and *5xFAD/Cnp^-/-^* mice ^73^. P-values from two-sided Student’s *t*-tests are shown.

Enrichment analysis revealed distinct patterns across the different disease conditions (Figure 5b). The pronounced amyloid-associated microgliosis in APPPS1 mice was characterized by strong enrichment of P1 and P2, whereas induction of the GPNMB-associated P3 program remained comparatively limited. In contrast, with increasing neuronal injury, P1 and particularly P2 enrichment declined, while P3 progressively increased and became most prominent in *Npc1^-/-^* and 4-month-old *DKO* mice. Analysis of the individual genes underlying these signatures further illustrated this differential regulation, with increasing expression of P3 genes across conditions with higher neuronal injury (Figure 5c). Importantly, mean P3 enrichment across the different conditions strongly correlated with plasma NfL levels (Pearson r = 0.88, p = 0.0098; Figure 5d), linking the GPNMB-associated transcriptional program to the extent of neuronal injury.

To independently validate the association between the Gpnmb-associated P3 program and increasing neuronal injury, we next asked whether a similar relationship is evident in an independent disease context combining myelin and Aβ pathology. Given the high degradative and lysosomal demand imposed on microglia by myelin turnover, and the contribution of lysosomal dysfunction to demyelination and impaired remyelination ^71,72^, we analyzed bulk RNA-seq data from *Cnp*^-/-^ mice with myelination deficits, alone or in combination with Aβ pathology in the *5xFAD* model ^73^(Figure 5e, f). P3 enrichment was increased in both *5xFAD* and *Cnp*^-/-^ mice but was most pronounced in *5xFAD Cnp*^-/-^ mice combining Aβ and myelin pathology (Figure 5e). This increase was reflected by individual P3-associated genes, including *Gpnmb*, *Spp1*, *Lgals3*, and *Apoe*, which showed highest expression under combined pathology (Figure 5f). Notably, myelin pathology alone was sufficient to induce several components of the P3 program, including *Gpnmb* and *Spp1*. In contrast, *Trem2* was strongly induced in 5xFAD mice but did not further increase to the same extent under combined pathology, while GPNMB-associated P3 genes did. This indicates that broad DAM activation can occur without a corresponding increase in the GPNMB-associated lysosomal response. Notably, the stronger P3 induction in *5xFAD* compared to *APPPS1* mice may reflect more pronounced motor phenotypes in *5xFAD* mice.

Together, these findings indicate that the composition of the microglial response changes with extent of neurodegeneration. While P1 and P2 programs are prominent in the *APPPS1* Aβ model, increasing neuronal injury is associated with a shift toward the GPNMB-associated lysosomal P3 program. Thus, broad canonical DAM activation and the GPNMB-associated lysosomal response can be dissociated, with the GPNMB-associated program more closely tracking the extent of neurodegeneration burden across disease contexts.

### The GPNMB-associated program is induced across lysosomal neurodegenerative disease models

Given the strong P3 enrichment observed in *Npc1^-/-^* mice in the cross-model transcriptomic analysis, we next examined the GPNMB-associated response and underlying lysosomal pathology in this model in greater detail. Consistent with substantial neuronal injury, plasma NfL levels were increased in *Npc1^-/-^* mice (Figure S6a).

DAM marker genes, including *Gpnmb, Spp1,* and *Clec7a* were markedly induced in the nCounter(R) GLIA-Panel analysis of whole brain RNA *from Npc1^-/-^* mice, yet the magnitude of induction was lower than in *DKO* mice, except for *Lgals3* and *Trem2* (Figure S6b). Thus, lysosomal/lipid trafficking impairment in *Npc1^-/-^* mice is accompanied by induction of a DAM transcriptional state that includes GPNMB, supporting the broader relevance of this response to lysosomal stress and degradative demand.

To assess regional patterns of GPNMB⁺ DAM in *Npc1^-/-^* mice, we performed immunofluorescence on sagittal brain sections. This revealed a regional specificity in GPNMB induction in microglia, especially in the cerebellum in regions where loss of Purkinje-cells has been described (Figure S6c) ^53^. Motivated by this regional induction, we dissected the brain and analyzed the hindbrain for microglia activation markers, and observed a pronounced elevation of GALECTIN3, CLEC7A, TREM2, and GPNMB (Figure S6d, e). Further, the hindbrain analysis revealed a significant increase in LC3 accompanied by the accumulation of RIPA-insoluble material in the UREA fraction, with clear deposition of p62 and ubiquitinated proteins. Strikingly, MBP was strongly reduced in *Npc1^-/-^* hindbrain (Figure S6d, e), a hallmark of NPC pathology ^53^ linking regional GPNMB⁺ DAM activation to areas of prominent tissue injury and myelin impairment.

We next expanded this analysis to *Ctsd^-/-^* (Cathepsin D deficiency), a second mouse model of lysosomal dysfunction and neurodegeneration ^50,74^. At postnatal day 28, we observed elevated levels of plasma NfL (Figure S6f), which were accompanied by a significant induction of GPNMB, GALECTIN3, CLEC7A and TREM2 in immunoblotting of total brain lysates (Figure S6g, h). The significant increase in LC3 was accompanied by a clear deposition of p62 and ubiquitinated proteins in the UREA fraction (Figure S6g, h) consistent with pronounced impairment in degradative capacity in *Ctsd^-/-^* mice. Together with the findings in *Npc1^-/-^*mice, these results support the conclusion that degradative overload and neurodegeneration are accompanied by induction of a GPNMB⁺ DAM program across distinct lysosomal disease contexts.

Collectively, analysis across *DKO*, *Npc1^-/-^* and *Ctsd^-/-^*mice revealed a convergent induction of GPNMB⁺ DAM under conditions of lysosomal impairment and degradative overload, with regional GPNMB accumulation associated with areas of pronounced neurodegenerative pathology.

### TREM2, but not CLEC7A, regulates DAM activation and induction of the GPNMB–associated program in *APPPS1* mice

Having established that the GPNMB-associated P3 program is associated with the extent of neuronal injury across disease models, we next asked whether its induction is linked to established upstream regulators of DAM activation. Because TREM2 is well established as a regulator of the microglial response to Aβ pathology, we compared *APPPS1* mice lacking *Trem2* or *Clec7a* ^2,5,75–77^. This provided a defined genetic context to determine whether the GPNMB-associated response depends on TREM2 and whether CLEC7A is similarly required.

The cross-model comparison (Figure 5) already indicated robust DAM activation in 16-month-old *APPPS1* mice, with *Trem2* and *Clec7a* robustly induced at the mRNA level, while *Gpnmb*, *Spp1* and *Lgals3* were also increased but remained markedly lower than in *DKO* mice (Figure S7b). Thus, *APPPS1* mice display strong DAM induction, but comparatively limited engagement of the GPNMB-associated P3 program.

Despite this lower baseline induction, loss of *Trem2* markedly attenuated the P3 response in *APPPS1* mice (Figure S7a, b). To examine this dependence across the broader Aβ-induced transcriptional response, we ranked genes according to their fold-change in *APPPS1* mice relative to WT controls (Figure S7c). Genes induced in *APPPS1* mice, including *Gpnmb*, *Spp1*, *Clec7a* and *Lgals3*, showed markedly reduced induction in *APPPS1 x Trem2*^-/-^ mice. In contrast, loss of *Clec7a* largely preserved the *APPPS1*-induced transcriptional response, including induction of GPNMB-associated genes. These findings indicate that, although P3 activation in *APPPS1* mice is weaker than in *DKO* mice, the GPNMB-associated response to Aβ pathology is strongly TREM2-dependent.

This differential dependence was further validated at the protein level (Figure S7d, e). *APPPS1* mice showed increased abundance of TREM2, CLEC7A, GPNMB and GALECTIN3, consistent with the transcriptional changes. Loss of TREM2 strongly reduced induction of CLEC7A, GPNMB and GALECTIN3, whereas loss of CLEC7A did not reduce TREM2, GALECTIN3 or GPNMB. GPNMB protein abundance was instead further increased in *APPPS1 x Clec7a*^-/-^ mice, indicating that CLEC7A is not required for induction of the GPNMB-associated response. Immunofluorescence analysis further confirmed accumulation of GPNMB⁺ IBA1⁺ microglia around Aβ pathology in *APPPS1* mice, with a marked reduction following loss of TREM2 but preservation in *APPPS1 x Clec7a*^-/-^ mice (Figure S7g).

Despite these pronounced differences in microglial responses, plasma NfL levels did not significantly differ between *APPPS1* mice and the corresponding *Trem2*-or *Clec7a*-deficient *APPPS1* groups at this age (Figure S7f). Together with the comparatively low NfL levels of *APPPS1* mice in the cross-model analysis (Figure 5a), this is consistent with the limited induction of the GPNMB-associated P3 program relative to *DKO* and other models with greater extent of neurodegeneration.

Together, these findings demonstrate that the GPNMB-associated program in *APPPS1* forms a part of the broader TREM2-dependent DAM continuum, whereas CLEC7A is dispensable for this induction.

### Loss of TREM2 is associated with exacerbated neurodegeneration under lysosomal stress

Across disease models, the GPNMB-associated P3 program was preferentially engaged under conditions of pronounced lysosomal burden and neurodegeneration. However, its association with increasing neuronal injury does not necessarily imply a detrimental function. Our CSF1R inhibition experiments demonstrated that attenuating microgliosis under lysosomal stress exacerbates neurodegeneration (Figure 1), suggesting a protective contribution of microglial responses in this context. We therefore investigated whether TREM2 loss attenuates the GPNMB-associated response and is associated with aggravated neuronal injury under lysosomal stress. To address this question, we examined aged *Grn^-/-^* mice with or without TREM2 deficiency. Given the breeding complexity associated with combining *Grn*, *Tmem106b*, and *Trem2* deficiencies, we used this established model of PGRN-associated lysosomal dysfunction to investigate the relationship between TREM2-dependent microglial responses and neurodegeneration. Previous analyses at 12 months of age showed that TREM2 deficiency did not rescue lysosomal pathology or the relatively modest neurodegenerative phenotype in *Grn^-/-^* mice ^78^. We therefore extended these analyses to 14-month-old mice to assess how TREM2 loss affects the GPNMB-associated program and neuronal injury at a more advanced disease stage. Immunoblotting revealed a prominent increase in GPNMB expression in *Grn^-/-^* animals, which was significantly reduced in *Grn^-/-^ x Trem2^-/-^* mice and absent in *Trem2^-/-^* mice (Figure S8a, b). This reduction was accompanied by a further increase in plasma NfL levels in *Grn^-/-^ x Trem2^-/-^* mice (Figure S8c), extending our previous findings of elevated CSF NfL in 12-month-old mice (Figure S8d) indicating aggravated neuronal injury following TREM2 loss (Figure S8c, d). Consistent with our findings in *APPPS1* mice, induction of GPNMB, GALECTIN3 and CLEC7A in *Grn^-/-^* mice is dependent on TREM2 signaling, while homeostatic ABI3 protein levels remained unaffected, indicating no change in homeostatic microglia (Figure S8a, b).

We next asked whether this relationship generalizes beyond PGRN-dependent pathology. A beneficial effect of TREM2-dependent microglia differentiation has been shown in TAU-P301L transgenic mice that were crossed to the PS2APP Aβ model and combined with a TREM2 loss of function ^79^. Enhanced neurodegeneration was described for TAU mice crossed to *Trem2^-/-^* mice. To obtain information on the transcriptional programs identified along the GPNMB gradient, we re-analyzed the published data ^79^. Analysis of P1, P2 and P3 signature enrichment across these models revealed a TREM2-dependent reduction of the GPNMB-associated P3 program in the context of increasing atrophy (Figure S8e). This further linked impaired induction of the P3 program with increased neurodegeneration burden following loss of TREM2 signaling.

Additionally, we investigated whether TREM2 agonism further enhances the GPNMB-associated program. Lengerich et al. showed that TREM2 agonism using a brain penetrant TREM2 antibody, ATV:4D9, in an Aβ mouse model (APP SAA) enhances mitochondrial metabolism, microglia activity, and glucose metabolism ^24^. We re-analyzed the published scRNA-seq data using cluster enrichment scores and observed that TREM2 stimulation increased the GPNMB-associated P3 program (Figure S8f), providing further evidence that TREM2 signaling regulates the GPNMB-associated response, consistent with a neuroprotective function of TREM2.

Together, these cross-model analyses provide further evidence that TREM2-signaling regulates induction of the GPNMB-associated program in disease-associated microglia. Across independent disease contexts, impaired TREM2-dependent induction of the GPNMB program was associated with greater neurodegeneration. Together, these findings support the GPNMB-associated program as a TREM2-regulated adaptive response that increases with lysosomal stress and ensuing neurodegeneration.

### The GPNMB-associated program is reversible upon rescue of lysosomal dysfunction in *DKO* mice

The tight coupling of the GPNMB-associated P3 program with lysosomal burden and neurodegeneration suggests an adaptive program engaged under degradative stress. We therefore asked whether alleviating lysosomal dysfunction is sufficient to reverse this GPNMB-associated program *in vivo* by leveraging the peripheral AAV-PGRN delivery, which rescues many features of FTLD-like pathology in *DKO* mice ^54^.

To understand how the transcriptional programs identified along the GPNMB gradient respond to lysosomal rescue, we re-analyzed bulk RNA-sequencing data from total brain tissue (Figure 6a-c). AAV-PGRN treatment markedly reduced enrichment of the GPNMB-associated P3 program in *DKO* mice, consistent with its close association with lysosomal dysfunction. P2 enrichment was also reduced following PGRN rescue, indicating a broader attenuation of DAM activation as pathology improved, whereas P1 showed a more limited response (Figure 6a, b). Consistent with the reduction of P3, *Gpnmb, Spp1* and *Lgals3*, together with the DAM-associated genes *Apoe, Clec7a* and *Trem2*, were strongly reduced toward WT levels following AAV-PGRN treatment (Figure 6c).

**Fig. 6:**
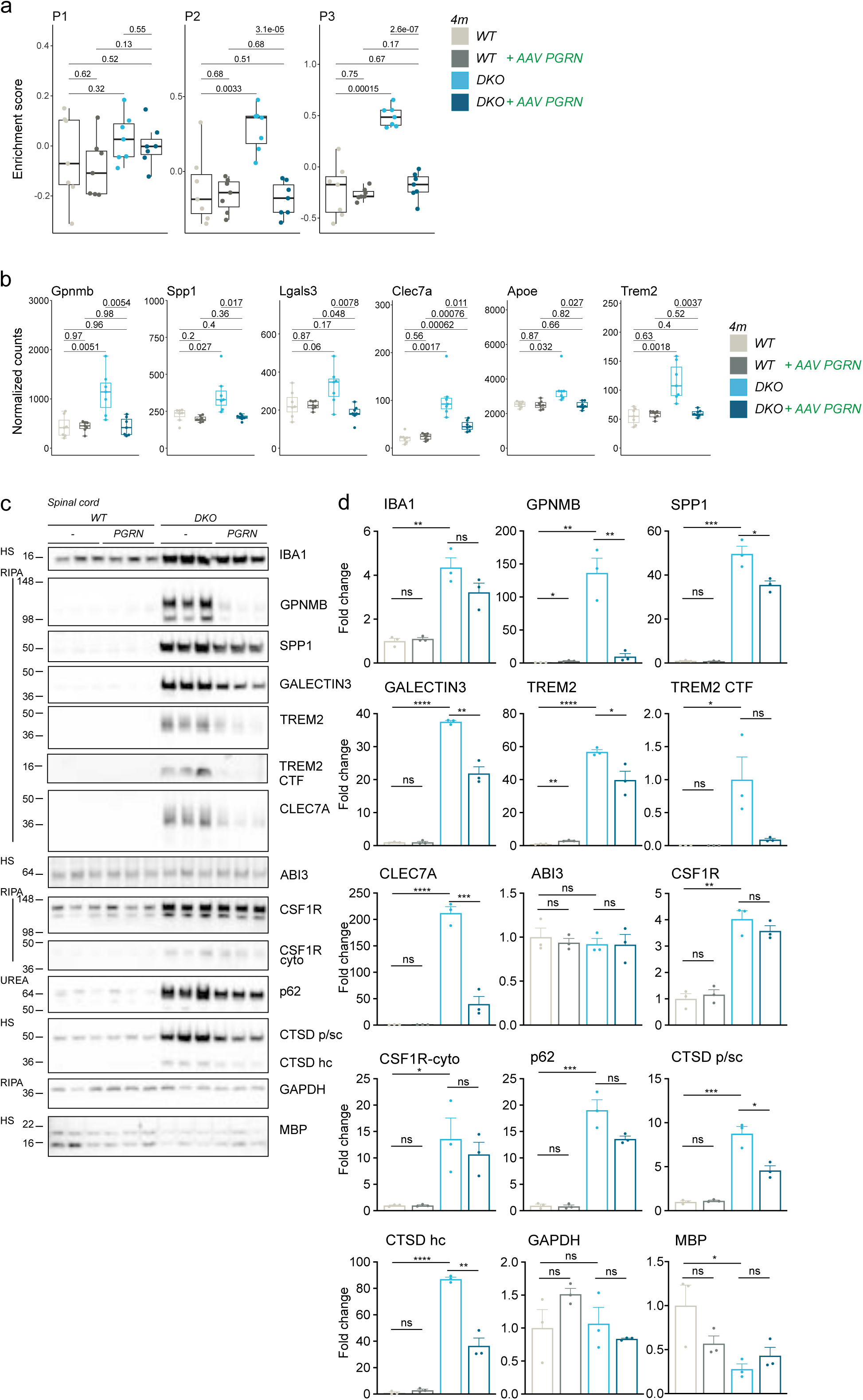
Lysosomal rescue reverses the GPNMB-associated program in *DKO* mice. (a) Bulk RNA-Seq data re-analyzed from ^54^. Total brain bulk transcriptomes of 4-month-old *WT* and *DKO* mice ± AAV PGRN treatment. Box plots showing enrichment values of filtered microglia signatures for clusters 1, 2, and 3, as depicted in (A) of 4-month-old *WT* and *DKO* mice ± AAV PGRN treatment. P-values from two-sided Student’s *t*-tests are shown. (b) Box plots of normalized counts of selected microglia genes in 4-month-old *WT* and *DKO* mice ± AAV PGRN treatment. P-values from two-sided Student’s *t*-tests are shown. (c) Western blot analysis of IBA1, GPNMB, SPP1, GALECTIN3, TREM2, CLEC7A, ABI3, CSF1R, p62, CTSD and GAPDH in the indicated fractions of spinal cord lysates from of 4-month-old *WT* and *DKO* mice ± AAV PGRN treatment, (n = 3 mice per group). Asterisk indicates unspecific band. (d) Quantification of proteins in (D), fold change to *WT* (n = 3 mice per group).

Dissection analysis of *DKO mice* revealed the strongest FTLD-like pathology in the hindbrain and the spinal cord (Figure S5a-c, 6c, d). Hence, immunoblotting revealed a significant reduction in GPNMB, SPP1, GALECTIN3, TREM2 and CLEC7A levels, together with partial normalization of CTSD and a tendency toward restoration of p62 and MBP following AAV-PGRN treatment (Figure 6c, d). Importantly, the accumulation of TREM2 CTF in the *DKO* was reduced following AAV-PGRN treatment, consistent with partial normalization of TREM2 processing (Figure 6c, d).

Collectively, these data show that correction of the underlying lysosomal defect by AAV-PGRN is accompanied by a pronounced reduction of the GPNMB-associated P3 program together with broader normalization of DAM responses. The reversibility of P3 supports the interpretation that the GPNMB-associated lysosomal program is dynamically engaged in response to lysosomal overload and associated tissue stress, rather than representing a fixed DAM state.

### Convergent lysosomal CSF signature tracks microglial activation and neuronal injury across neurodegeneration mouse models

Given the lysosomal overload associated with the GPNMB-associated microglial program, we asked whether these changes are reflected in a detectable CSF proteomic signature. We performed mass spectrometry-based proteomics on cerebrospinal fluid (CSF) from *DKO* and *WT* mice (Figure 7a). We detected peptides from 1467 proteins of high quality in the CSF. Among these, 672 proteins were significantly elevated in the *DKO* CSF and a subset of 79 proteins overlapped with the differentially expressed genes from the scRNA-seq analysis, linking the CSF profile to the DAM transcriptional phenotype (Figure 7a, b, Suppl. Table 6). Prominently elevated proteins included microglia-associated proteins such as GPNMB, TREM2, C1QA, LYZ2, APOE, AXL, LAG3 and SPP1. David pathway analysis revealed a high abundance of lysosomal proteins, together with proteins associated with neuronal and axonal function. The robust detection of lysosomal and neuronal proteins in the CSF of *DKO* mice prompted us to assess the human relevance. We compared these CSF changes to published proteomic data sets derived from CSF of FTD patients ^60,80^. Of the 66 proteins reported as altered in FTD patients, 32 were significantly elevated in *DKO* mouse CSF analysis (Figure 7b), indicating substantial overlap with human pathology.

**Fig. 7:**
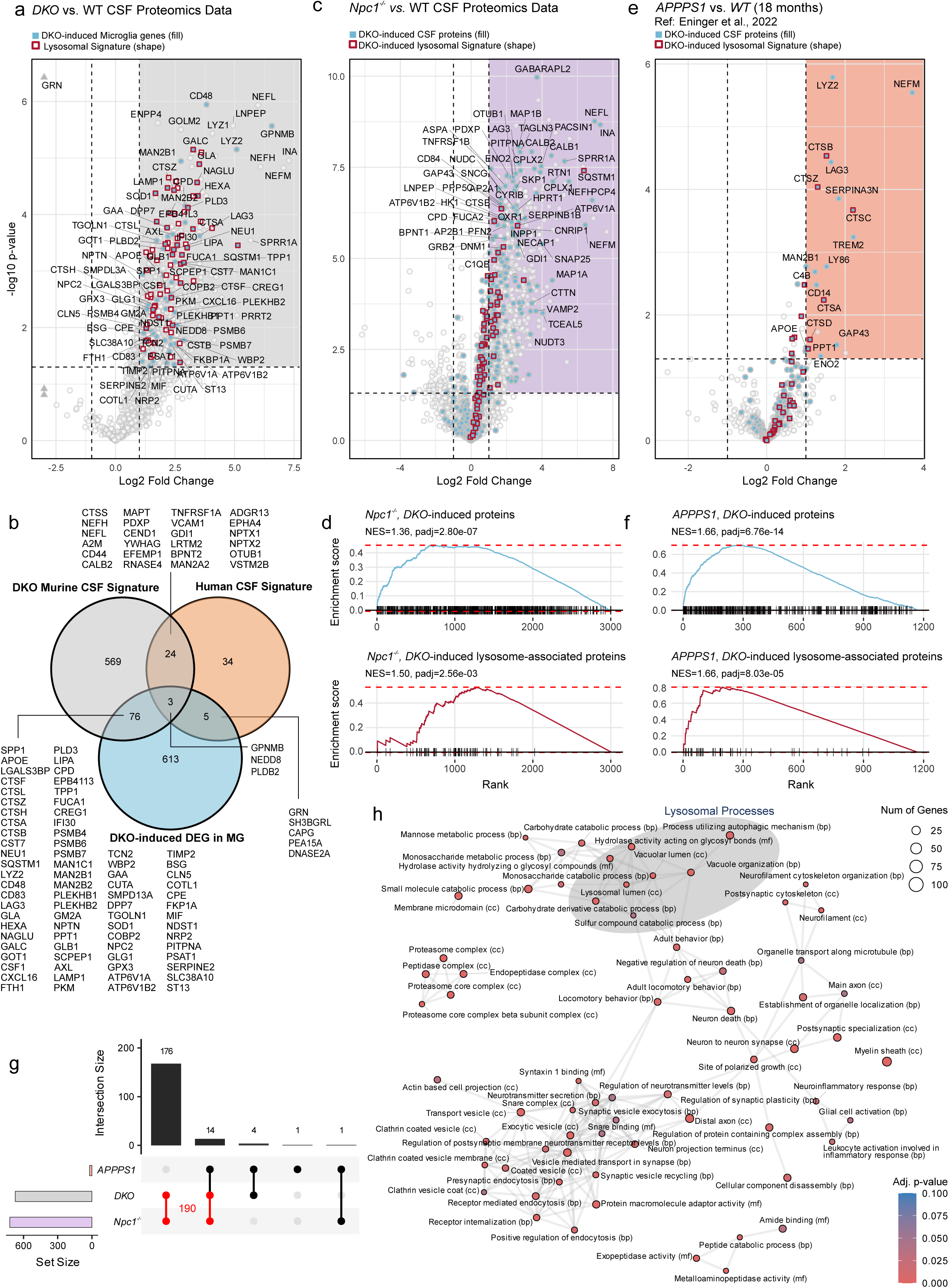
A convergent lysosomal CSF signature reflects microglial activation and neuronal injury. (a) Volcano plot showing –log(10) p-values and log(2) fold changes of proteins detected by proteomic analysis in CSF from 4-month-old *DKO* vs *WT* mice (n = 4 per group). Proteins overlapping with enriched signatures related to lysosomal activity are outlined in red, while those overlapping *DKO*-induced differentially expressed genes in microglia from scRNA-seq data (Fig. 2) are filled in blue. For visualization, fold changes below −3 were capped and denoted by triangle symbols. (b) Venn diagram showing overlaps between *DKO*-induced proteins in CSF from 4-month-old *DKO* mice compared to *WT* controls, human CSF protein signatures from patients with neurodegenerative diseases ^60,80^, and *DKO*-induced differentially expressed genes in microglia from scRNA-seq data (Fig. 2). (c) Volcano plot showing –log(10) p-values and log(2) fold changes of proteins detected by proteomic analysis in CSF of 2-month-old *WT* and *Npc1^-/-^* (n = 5 per group). CSF proteins overlapping with *DKO*-induced differentially expressed proteins are shown in blue (Fig. 7G), whereas proteins additionally overlapping lysosome-related enriched signatures are highlighted with a red outline. (d) Ranked enrichment of *DKO*-induced CSF-signature proteins, in *Npc1^-/-^* CSF in blue. Ranked enrichment of *DKO*-induced lysosome associated CSF-signature proteins, in *Npc1^-/-^*CSF in red. (e) Reanalysis of CSF proteomics data from at 18-month-old *WT* and *APPPS1* mice ^81^. Volcano plot showing –log(10) p-values and log(2) fold changes of proteins in *APPPS* vs. *WT* mice. Proteins overlapping *DKO*-induced differentially expressed proteins from CSF proteomics (Fig. 7G) are shown in blue, whereas proteins additionally overlapping lysosome-related enriched signatures are highlighted with a red outline. (f) Ranked enrichment of *DKO*-induced CSF-signature proteins, in *APPPS1* CSF in blue. Ranked enrichment of *DKO*-induced lysosome associated CSF-signature proteins, in *APPPS1* CSF in red. (g) Upset plot depicting CSF proteomics–based intersections of proteins significantly induced in *DKO, Npc1^-/-^, APPPS1*, and *A30P-aSyn* mice relative to their respective controls. Proteins detected in both *DKO* and *Npc1*^-/-^ are highlighted in red. (h) Network visualization of Gene Ontology enrichment results based on CSF-signature of 190 proteins found to be elevated *DKO* and *Npc1^-/-^* mice. Colored according to adjusted p-values and scaled according to the number of proteins enriched per term. Edges indicate overlapping genes.

To test whether this CSF lysosomal signature can be generalized and validated in an independent neurodegeneration model, we performed CSF proteomics in *Npc1^-/-^* and *WT* mice. We detected peptides from 3006 proteins with 720 proteins significantly elevated in *Npc1^-/-^*CSF (Figure 7c, Suppl. Table 7). Ranked enrichment analysis confirmed a robust overlap with the proteins that were also induced in the *DKO* mice, with a high abundance of lysosomal, autophagy-associated, proteasomal and axonal/filament related proteins (Figure 7d), including the shared induction of microglia-associated proteins such as GPNMB, TREM2, ATP6V1A, C1QA, LAG3 and SPP1 (Suppl. Table 8). Consistent with the modest neurodegeneration in Aβ models relative to lysosomal impaired neurodegeneration models, published CSF datasets from *APPPS1* mice at an age of 18 months revealed a weaker, yet directionally similar, lysosomal signature (Figure 7e, f) ^81^. Notably, the overlap with proteins induced in the *DKO* model was highly significant in both systems, further supporting a convergent CSF signature associated with lysosomal dysfunction across neurodegeneration models.

To further refine the CSF signature associated with lysosomal dysfunction, we focused on the overlap between the two lysosomal impaired neurodegeneration models *DKO* and *Npc1^-/-^*. This analysis identified a shared core set of 190 proteins that were significantly induced in the CSF of both models (Figure 7g, Suppl. Table 8). These proteins are particularly informative for lysosomal dysfunction as a convergent mechanistic axis in neurodegeneration and partially overlapped with proteins detectable in the CSF of *APPPS1,* supporting broader relevance across disease etiologies. GO enrichment analysis of the 190-protein core set confirmed strong overrepresentation of lysosome-, autophagy- and proteasome-related pathways but also revealed functional links between lysosomal dysfunction and neurodegeneration-relevant processes including locomotion, glial cell activation, and immune response regulation (Figure 7h).

Together, these data support a model in which the GPNMB-associated microglial program becomes increasingly engaged under high degradative demand and is accompanied by coordinated lysosomal and lipid-handling responses.

## Discussion

We identify a GPNMB-associated lysosomal program within the broader DAM response that is consistently engaged under sustained lysosomal overload across neurodegeneration models, representing a dynamic microglial adaptation to increased degradative demand early in disease that likely enhances clearance capacity and limits neuronal damage. However, with sustained lysosomal burden, increasing engagement of this program coincides with lysosomal substrate accumulation, crystal formation, and worsening neurodegeneration. In this framework, the GPNMB-associated program is not a fixed endpoint but an inducible and, importantly, reversible component of the DAM response, whose magnitude tracks lysosomal stress and extent of neurodegeneration.

Although GPNMB⁺ cells have been linked to detrimental phenotypes in cancer, where GPNMB has been described as a senescence-specific antigen that promotes survival of senescent cells, immunosuppression and anti-inflammation ^82,83^, our data support a different role in neurodegeneration, where increasing GPNMB expression within DAM is associated with a predominantly lysosomal and lipid-metabolic transcriptional profile, consistent with a microglial program engaged under high degradative demand. Importantly, ordering DAM according to increasing Gpnmb expression showed that it marks progressive remodeling within the broader DAM spectrum. As the GPNMB-associated lysosomal program increases, inflammatory and chemokine-related, as well as endosomal programs become less prominent, while lysosomal and lipid-handling genes are progressively induced. We propose that the GPNMB-associated program is induced by neuronal injury and/or as a compensatory response to impaired lysosomal integrity, reflecting a phagocytic, microglial clearance program that is also engaged during physiological remodeling, including development and age-associated white matter turnover ^84–86^. In neurodegeneration, specific PU.1^low^ DAM have been shown to adopt distinct protective functions that restrain neuroinflammation and thereby mitigate AD progression ^87^. Thus, rather than implying a consistently detrimental state, the GPNMB-associated program may reflect an initially adaptive microglial response that supports tissue maintenance under degradative stress, even if chronic lysosomal overload may constrain its functional capacity.

Microglial activation is increasingly recognized as heterogeneous and context-dependent ^88^. We set out to identify a program specific to neurodegeneration and shared across mouse models. Using scRNA-seq and orthogonal validation, we identified a microglia cluster expanded in *DKO* mice, characterized by elevated levels of *Gpnmb, Spp1, Lpl, Igf1*, and *Fabp5* and enrichment of lysosomal and lipid metabolism genes. This cluster partially overlaps with previously described DAM genes, yet is distinguished by a pronounced lysosomal signature and reduced sensitivity to CSF1R inhibition. Spatial analysis further showed that GPNMB⁺ microglia were particularly abundant in regions exhibiting neurodegeneration, myelin loss and enriched for neuropathological hallmarks such as p62, phosphorylated TDP-43, and accumulation of ubiquitinated proteins. Electron microscopy revealed crystal accumulation within lysosomal compartments of microglia close to neuritic dystrophy, consistent with cholesterol crystallization described in the context of lipid overload ^66,67,69^. Notably, induction of the GPNMB-associated program represents a convergent response under high lipid and protein clearance demand and tracks the extent of neurodegeneration and lysosomal pathology across several mouse models (e.g. *Npc1^-/-^* and *Ctsd^-/-^* mice).

A central insight gained from cross-model comparisons is that Aβ models robustly activate DAM but show comparatively limited neurodegeneration ^55^. Consistent with the lack of strong plasma NfL elevation, the GPNMB-associated P3 program was comparatively modestly induced in aged APPPS1 mice. Notably, *APPPS1* mice showed strong enrichment of P1 and P2 despite comparatively limited P3 induction, indicating that pronounced plaque-associated DAM activation can occur without strong engagement of the GPNMB-associated lysosomal program. Re-analysis of the ^73^data further showed that P3 enrichment was increased by both Aβ and myelin pathology and was most pronounced when these pathologies were combined, identifying myelin pathology as a context associated with enhanced induction of the GPNMB-associated program. Consistent with our framework, a recent human myeloid single cell atlas identified a GPNMB-high disease associated subtype that expands with AD pathology and exhibits neuroprotective activity ^49^. Additionally, human iPSC-derived microglia can be driven toward a GPNMB⁺ DAM transcriptional state by exposure to pathological substrates, including Aβ and apoptotic neurons ^89^.

Together, these comparisons suggest a tipping point model in which the GPNMB-associated program emerges as degradative demand approaches or exceeds lysosomal capacity or reserve. The GPNMB-associated program may contribute to lysosomal stress management, by potentially modulating the routing of autophagy-associated cargo toward lysosomal degradation under sustained load to help preserve residual lysosomal capacity within the GPNMB⁺ DAM state. In lysosome-driven models (*DKO*, *Npc1^-/-^* and *Ctsd^-/-^*), the GPNMB-associated program is particularly prominent and coincides with marked lysosomal pathology and neurodegeneration, whereas in Aβ models its induction is more limited but increases when chronic plaque burden converges with myelin disruption and impaired lipid handling. Conceptually, the onset and magnitude of the GPNMB-associated program, together with the CSF lysosomal signature, may therefore mark the point at which microglia transition from effective compensation toward overload and declining clearance capacity, potentially accelerating neurodegeneration. Assessing GPNMB-associated markers may provide a framework to assess disease progression beyond pathology burden alone. Monitoring when the GPNMB-associated lysosomal program becomes strongly engaged could help define a therapeutic window in which restoring lysosomal capacity or modulating microglia is most likely to delay transition into irreversible neurodegenerative decline.

Our original attempt to reduce microgliosis and ameliorate the FTLD-like pathology revealed a general principle: Microglial interventions can have fundamentally different outcomes depending on disease context and stage. CSF1R-inhibition strongly reduced microglial abundance in *WT* mice without compromising neuronal integrity, but in the *DKO* model, where microglia are challenged by lysosomal dysfunction, CSF1R-inhibition exacerbated neurodegeneration. Similar context dependence has been observed in Aβ mouse models where microglia depletion initially reduced plaque load and associated toxicity ^9,90,91^. However, prolonged microglial ablation exacerbates plaque-associated pathology, including increased dystrophic neurites surrounding fibrillar Aβ deposits. The complete loss of microglia in 5xFAD FIRE mice led to increased vascular Aβ and early lethality ^12,92^. A possible explanation is that lysosomal burden eventually exceeds the microglial capacity required to cope with lysosomal stress during tissue maintenance. Although GPNMB expression has been associated with reduced inflammatory capacity, enhanced phagocytosis and autophagy ^93,94^, our data indicate that partial persistence of the GPNMB⁺ DAM state under CSF1R-inhibition is not sufficient to prevent neuronal loss in the *DKO* setting. Notably, while GPNMB protein levels remained elevated, the abundance of GPNMB⁺ DAM declined, implying that the remaining cells may further upregulate GPNMB as a compensatory response, yet cannot fully sustain clearance demands. This imbalance is consistent with a scenario in which the GPNMB-associated program reflects an attempted protective program that becomes functionally constrained when microglial numbers are depleted under conditions of high degradative stress. Additionally, GPNMB high expressing cells showed reduced expression of immune-sensing, endosomal trafficking and motility programs and increased expression of factors that may restrain extracellular proteolysis and tissue remodeling (e.g. CST7), consistent with a shift toward a more tissue-preserving response.

To further investigate regulation of the GPNMB-response and its relationship to TREM2-dependent neuroprotection, we leveraged TREM2 loss-of-function, given the established role of TREM2 in DAM induction during Aβ pathology, lipid sensing, plaque engagement, and metabolic fitness ^2,5,7,21^. In aged *APPPS1* mice, the modest GPNMB-associated response was reduced by TREM2 deficiency, placing this program within the broader TREM2-dependent response to Aβ pathology, whereas CLEC7A deficiency did not measurably alter the microglial response to Aβ, consistent with CLEC7A serving as an activation marker rather than a required regulator in this setting ^77^. We note that TREM2 broadly reshapes the microglial response to Aβ, which would require time-resolved perturbation studies to distinguish direct regulation of the GPNMB-associated program from broader effects on DAM development.

Extending beyond amyloid, TREM2 loss in *Grn^-/-^* mice markedly suppressed the GPNMB-associated response and was associated with increased NfL, consistent with a potentially protective role of the broader TREM2-dependent microglial response under lysosomal stress. Given the abundant microglial expression of PGRN and its partial lysosomal localization, this finding is particularly relevant because aged *Grn^-/-^* mice exhibit neuropathology and elevated DAM gene expression ^44,46,95–98^

Further support comes from published TAU-P301L and Aβ combination models showing aggravated neurodegeneration with reduced induction of the GPNMB-associated P3 program upon TREM2 loss ^79^. Conversely, application of a TREM2-stimulating antibody (ATV:4D9) increased the GPNMB-associated P3 program ^24^ Together, our data suggest that increasing GPNMB expression within DAM is accompanied by lysosomal and lipid-handling programs but can simultaneously exhibit signs of impaired lysosomal processing. This is consistent with compensatory induction of the lysosomal machinery that becomes insufficient under chronic overload. GPNMB itself localizes to endosomal and lysosomal compartments and has been implicated in regulating phagocytosis, lysosomal integrity, and autophagy coupling to lysosomes ^44,47,48,99,100^.

Together, these observations support a model in which microglia adopt a metabolically and epigenetically reprogrammed response to cope with intracellular degradative burden. A key feature of our framework is reversibility, supporting the view that the GPNMB-associated lysosomal program is dynamically regulated rather than representing a terminal microglial fate. Peripheral AAV-PGRN delivery in *DKO* mice (Reich et al. 2024) rescued features of lysosomal dysfunction and markedly reduced the GPNMB-associated P3 program. This highlights that the GPNMB-associated response is dynamically coupled to the underlying lysosomal and pathological burden and can recede when this burden is alleviated.

Finally, CSF proteomics detected numerous lysosomal and microglial proteins overlapping with the DEG microglial signature in *DKO* mice including GPNMB, SPP1, CTSB, CTSS, CTSL, and LYZ2. Many elevated proteins were of microglial and neuronal origin and 32 proteins overlapped with those previously reported to be elevated in the CSF of human AD and FTD patients ^60,80,101–105^. The overlap with induced proteins in *Npc1^-/-^* CSF samples and a weaker induction in Aβ contexts suggests that CSF profiles may capture convergent lysosomal and microglial responses and provide potential translational readouts of lysosomal dysfunction and neurodegeneration. In parallel, soluble GPNMB has been proposed as a biomarker in lysosomal storage and neurodegenerative disorders ^97,106–111^, aligning with our observation that GPNMB tracks lysosomal stress across models.

This study is based on a variety of murine models, that capture key features of neurodegeneration, inflammation and lysosomal dysfunction, but may not reflect the full diversity of human microglia states. Moreover, much of the cross-model analysis is correlative, and perturbation of TREM2 broadly affects DAM activation, preventing the effects of TREM2 loss from being fully attributed to the GPNMB-associated program. Further perturbation studies of genes highly expressed within the GPNMB-associated program will be important to define their functional contributions, because GPNMB serves as a marker of this microglial state rather than its driver. Such experiments will require particular caution, as further compromising components of an adaptive lysosomal response under conditions of impaired degradative capacity may exacerbate disease progression.

Nevertheless, the strong cross-model consistency and the substantial overlap of transcriptional, epigenetic, biochemical and CSF proteomic evidence support the interpretation of the GPNMB-associated program as a dynamic microglial response to lysosomal stress and the extent of neurodegeneration. The overlap of our CSF lysosomal signature with human FTLD datasets further supports the translational relevance of this response.

In summary, we identify a GPNMB-associated lysosomal program that is progressively engaged within the broader DAM continuum under conditions of lysosomal overload, neurodegeneration and myelin injury. Our data support a framework in which this program is initially adaptive to increasing degradative stress but may be increasingly constrained under chronic lysosomal overload. This refined framework identifies the TREM2-regulated GPNMB-associated program and its associated biomarkers as potential entry points for stratifying lysosomal stress and neurodegenerative severity across neurodegenerative diseases.

## Resource availability

Further information and requests should be directed to and will be fulfilled by Georg Werner and Joachim L. Schultze.

## Materials availability

A new CLEC7A antibody for immunoblotting was generated in this study and a new CLEC7A Knockout mouse line was established by CRISPR CAS9.

## Data and code availability

All single-cell transcriptomic data will be deposited at the Gene Expression Omnibus (GEO) under access numbers (Access to GEO repository via the token ). Original code will be deposited at Gitlab and is publicly available as of the date of publication. Additional processed transcriptomics data are available upon reasonable request.

The proteomics data will be deposited to the ProteomeXchange Consortium via the PRIDE partner repository with the dataset identifier (Token: , Username: Password: ). Processed data and analysis scripts are available upon reasonable request.

## Study overview

To identify a neurodegeneration specific microglia population and evaluate the contribution of microglial activation on neurodegeneration, a FTLD-like mouse *Grn^-/-^* /*Tmem106b^-/-^*(*DKO*) model was utilized. We performed TSPO-PET imaging, immunohistochemistry and single cell sequencing after pharmacological microglia depletion in *WT* and *DKO* mice. Analysis of *Npc1^-/-^* , *Ctsd^-/-^* and Aβ mouse models *APPPS1, APPPS1*x*Trem2^-/-^* and *APPPS1*x*Clec7a^-/-^*by bulk RNA analysis and biochemistry confirmed that GPNMB⁺ DAM correlate with severity of neurodegeneration, myelination impairment and lysosomal dysfunction.

## Animal Ethics Approval

All animal experiments were performed in accordance with the German animal welfare legislation and approved by the government of upper Bavaria (Regierung von Oberbayern) under the animal license: **ROB-55.2-2532.Vet_02-17-106**. Mice were kept under standard housing conditions including standard pellet food and water provided *ad libitum*. Mixed-sex cohorts were used unless stated.

## Method details

Detailed description of the methods applied is attached.

## Acknowledgements

We would like to thank Dr. M. Nishihara (Department of Veterinary Physiology, the University of Tokyo) for providing the *Grn ^-/-^* founder mouse strain. Markus Damme and Paul Saftig for providing *Ctsd ^-/-^*brain material. We thank Claudia Krenner, Katrin Buschmann, Cornelia Niemann, Pardis Khosravani for excellent technical assistance. The authors like to thank Michael Heide (Core Facility “Digital Single Molecule/NanoString Technologies”; Deutsches Konsortium für Translationale Krebsforschung, Partner Site München, Labor für Experimentelle Leukämie-und Lymphom-Forschung (ELLF)) for supporting NanoString measurements. We would like to acknowledge the NanoString Technologies EMEA grant program for providing a Glia panel.

## Author contributions

Conceptualization: GW, JSS, CH, AC, JLS

Methodology, Investigation: GW, RR, EDD, JSS, JG, GK, KN, LD, JCMS, SH, AFF, PH, BN, MR, IP, SP, AVC, SAM, MW, SL, CS, JBS, DP AR, SR, MJS, TL, BW, WW, RF, AL, DE, KK

Data analysis: GW, JSS, JG, KN, SFL Visualization: GW, JSS

Supervision: GW, MDB, AG, CG, GDP, ST, SFL, MSi, MSc, MB, CH, AC, JLS

Funding acquisition: CH, ST, JLS

Writing, original draft: GW, JSS, CH, AC, JLS Writing – Review & Editing: all authors

## Method details

Werner, Schulte-Schrepping, et al., 2026

### Microglia depletion

Pharmacological depletion of microglia by CSF1R inhibition was performed on 2-month-old male and female *DKO* and *WT* mice for a duration of 8 weeks with the CSF1R inhibitor PLX 5622 as established by ^1^ PLX5622 formulated chow was used at 1200 mg/kg (Medchempress, CAS No: 1303420-67-8), according to established protocols ^1^. *WT* and *DKO* cohorts received identical dosing regimens; control groups received matched chow without PLX5622. Food was renewed every 3 to 4 days.

### TSPO-µPET

Small animal positron emission tomography (µPET) procedures were acquired on a nanoScan PET/MR system (Mediso Ltd., Hungary) using a three-bed imaging chamber and followed a standardized protocol for radiochemistry, acquisition, and post-processing^2,3^ . TSPO µPET with a 60 to 90 min emission window was used to measure microglial activity, after injection of 12.2 ± 2.1 MBq [^18^F]GE180. In addition, two 2-min anatomical T1 weighted MR scans (axial and sagittal) were performed (head receive coil, matrix size 96 × 96 × 22, voxel size 0.24 mm × 0.24 mm × 0.80 mm, repetition time 677 ms, echo time 28.56 ms, flip angle 90°). The respective T1 images were used to create a body-air material map for the attenuation correction. µPET list-mode data were reconstructed using a 3D iterative algorithm (Tera-Tomo 3D, MedisoLtd, Hungary) with 8 iterations and 6 subsets using a matrix size of 283 x 283 x 313 voxels of 0.3 x 0.3 x 0.3 mm^3^. Random, scatter, attenuation, and decay corrections were applied. Male and female *DKO* and *WT* mice underwent longitudinal TSPO µPET at 2.0 and 3.5 months of age. PMOD (V3.5) was used to analyze the data. TSPO µPET SUV estimates from a brainstem target VOI (7 mm³) were compared descriptively between the two genotype groups by normalizing tissue activity to the injected dose adjusted for body weight (injected activity/mouse weight).

### Immunofluorescence staining

Following deep anesthesia, mice were PBS perfused. One brain hemisphere of each mice was fixed overnight in 4% paraformaldehyde (PFA) in PBS (pH = 7.4) and transferred to 30% sucrose in PBS. 35 μm thick free-floating sections were cut on a sliding microtome (Leica).

Sections were blocked in TBST 1x, 0.5 % TritonX-100, containing 1% donkey serum and incubated with primary antibodies overnight at 4°C. After 3 rounds of washing, fluorophore-conjugated secondary antibodies (Alexa Fluor Secondary, Thermo Fisher Scientific) were applied for 2 h. After additional washing steps, sections were embedded in Fluoromount^TM^ (Sigma-Aldrich). Images were taken on a LEICA DMI-8 with a DFC9000GT camera and objective: (HC PL FL L 20× / 0.40 CORR PH1). Confocal Images were acquired on Zeiss LSM 800 Microscope Axio Observer 7 (Objective Plan-Apochromat 63×/1,4 Oil M27). Quantification was performed in FIJI.

### Plasma neurofilament light chain (NfL) quantification

Plasma NfL was quantified using the Quanterix Simoa Neurofilament Light Advantage kit. Plasma samples were run in duplicates, at a 40x dilution, following the NfL analysis protocol on the Quanterix SR-X instrument.

### Isolation of glial cell populations

Following deep anesthesia, *WT* and *DKO* mice were PBS perfused and whole brains were dissociated using the Adult Brain Dissociation Kit (Miltenyi, 130-107-677). 3 animals per genotype and treatment were analyzed. Microglia, astrocytes, and oligodendrocytes were enriched together with the Neuron Isolation Kit (Miltenyi,130-115-390) containing CD11b, ACSA-2, and O4 MicroBeads, respectively. Living cells were sorted on a BD FACSAria^TM^ Fusion Flow Cytometer and immediately processed for scRNA-seq.

### Single-cell RNA sequencing using BD Rhapsody

ScRNA-seq of purified microglia, astrocytes, and oligodendrocytes was performed using the BD Rhapsody™ Express Single-Cell Analysis System. Single cells were isolated according to the manufacturer’s recommendations using the BD Rhapsody Cartridge Kit (BD Biosciences, cat. no. 633733) and the BD Rhapsody Cartridge Reagent Kit (BD Biosciences, cat. no. 633731). cDNA was generated using the BD Rhapsody cDNA Kit (BD Biosciences, cat. no. 633773). Whole transcriptome libraries were prepared using the BD Rhapsody Whole Transcriptome Analysis Amplification Kit (BD Biosciences, cat. no. 633801). Individual samples were run on separate cartridges across multiple experimental dates. Final libraries were quantified using a Qubit Fluorometer with the Qubit dsDNA HS Kit (Thermo Fisher Scientific), and fragment size distributions were assessed using the Agilent high-sensitivity D5000 assay on a TapeStation 4200 system (Agilent Technologies). Libraries were sequenced on an Illumina NovaSeq 6000 platform in paired-end mode (PE75), using NovaSeq 6000 S1 Reagent Kit (200 cycles) chemistry.

### Preprocessing, demultiplexing, and quality control

After demultiplexing of bcl files using bcl2fastq2 (v2.20) from Illumina and quality control, paired-end scRNA-seq reads were filtered for valid cell barcodes using the barcode whitelist provided by BD. Cutadapt 1.16 was then used to trim NexteraPE-PE adaptor sequences where needed and to filter reads for a PHRED score of 20 or above. Then, STAR 2.7.3a was used for alignment against GRCm39 reference genome and the gencode vM33 transcriptome annotation. Dropseq-tools 2.0.0 were used to quantify gene expression and collapse to UMI count data (https://github.com/broadinstitute/Drop-seq/).

Downstream analysis of BD Rhapsody scRNA-seq data was performed in R (v4.3.3). ScRNA-seq UMI count matrices were imported and gene expression data analysis was performed using the *Seurat* package (v5.0.3). Doublet detection was performed using DoubletFinder (v2.0.4) on cells from each experimental unit separately (PCs = 1:20, pN = 0.25, pK = 0.005) and only cells classified as singlets were retained for downstream analyses. Quality control filtering was applied to remove low-quality cells. Cells were excluded if they expressed fewer than 250 features or exhibited mitochondrial transcript content exceeding 25% of total counts.

### Normalization, variable feature selection, and dimensionality reduction

Gene expression values were normalized using Seurat’s *LogNormalize()* method by scaling total UMI counts per cell to 10,000 followed by log-transformation. Highly variable genes were identified using the variance-stabilizing transformation (vst) method, selecting the top 2,000 variable features.

The data were scaled and centered without regressing for any factors. Principal component analysis (PCA) was performed on the scaled expression matrix using the selected variable genes. For downstream analyses, the first 20 principal components were retained for batch correction, dimensionality reduction and clustering. Batch correction was performed using harmony (v1.2.0) based on the first 20 principal components setting experimental date as the batch variable. Uniform Manifold Approximation and Projection (UMAP) based on harmony corrected components was used for two-dimensional visualization based on the selected principal components.

### Clustering, cluster marker gene identification and cell type annotation

Cell clustering was performed using a shared nearest neighbor (SNN) graph constructed from the first 20 harmony-corrected principal components. Graph-based clustering was carried out using the Louvain algorithm implemented in Seurat with a resolution parameter of 0.2. Clusters were annotated manually based on the expression of known marker genes and label transfer using published murine brain scRNA-seq data sets^4,5^. Cluster-specific marker genes were identified using the Wilcoxon rank-sum test as implemented in Seurat’s *FindAllMarkers()* function. Differential expression of marker genes was assessed by comparing each cluster against all remaining cells. Genes were considered if they were expressed in at least 20% of cells within a cluster, showed a minimum log-fold change of 0.2, and exhibited a minimum difference of 10% in the fraction of expressing cells between groups.

### Differential gene expression analysis among experimental conditions

Differential gene expression analyses between experimental conditions (*WT*, *DKO*, *DKO*+PLX5622) across all microglia were performed using the Wilcoxon rank-sum test as implemented in Seurat *FindMarkers()*. Genes were considered statistically significantly different if they were expressed in at least 10% of cells in either group, exceeded a minimum log-fold change threshold of 0.25 and exhibited a minimum difference of 10% in the fraction of expressing cells between groups. P values were adjusted for multiple testing using the Bonferroni procedure, and adjusted p values < 0.1 were considered statistically significant.

### Subset analysis of microglia Microglia marker gene detection

For the specific analyses of microglia, the dataset was subsetted to include only cells from the clusters Microglia_1, Microglia_2, and Microglia_3. Microglia cluster-specific marker genes were identified using the Wilcoxon rank-sum test as implemented in Seurat’s *FindAllMarkers()* function. Differential expression of marker genes was assessed by comparing each cluster against all remaining cells. Genes were considered if they were expressed in at least 10% of cells within a cluster and showed a minimum log-fold change of 0.2. P values were adjusted for multiple testing using the Bonferroni procedure, and adjusted p values < 0.1 were considered statistically significant. The resulting lists of microglia cluster-specific marker genes were used for signature enrichments across independent data sets.

### Gene ontology enrichment analysis

Functional enrichment analyses were performed in R using clusterProfiler (v4.10.1) by applying over-representation testing with the *enricher()* function. Gene set definitions for Gene Ontology biological processes and cellular compartment were loaded from MSigDB ^6^GMT files (m5.go.bp.v2023.1.Mm.symbols.gmt, m5.go.cc.v2023.1.Mm.symbols.gmt). For each tested gene list, enrichment was computed against a background of present genes detected in the respective Seurat object subsetted per cluster, defined as genes with row-summed raw counts > 20. P values were adjusted using Bonferroni correction, and enrichment results were filtered using a qvalue cutoff of <0.1.

### GPNMB expression–ordered analysis of disease-associated microglia

To characterize transcriptional changes associated with increasing *Gpnmb* expression within DAM, cells from microglia cluster 3 were ordered according to single-cell *Gpnmb* expression and divided into discrete expression bins. Cells lacking detectable *Gpnmb* expression were separated from *Gpnmb*-expressing cells, and *Gpnmb*-positive cells were ranked according to expression level and distributed across ordered bins. Homeostatic microglia from cluster Microglia_1 were included as an additional reference population to relate transcriptional changes within cluster 3 to the transition from homeostatic to disease-associated microglia.

For differential expression analysis, cells were grouped into homeostatic microglia, GPNMB-low microglia, and GPNMB-high microglia. GPNMB-low and GPNMB-high cells comprised the lower and upper halves of the ordered *Gpnmb*-expressing bins, respectively. Differentially expressed genes were identified using Seurat FindMarkers() for three pairwise comparisons: GPNMB-low versus homeostatic microglia, GPNMB-high versus GPNMB-low microglia, and GPNMB-high versus homeostatic microglia. Genes were tested using a minimum fraction of expressing cells of 0.1, a log-fold-change threshold of 0.2, and a minimum difference in the fraction of expressing cells of 0.1. Genes with an adjusted *p*-value < 0.05 in at least one of the three comparisons were retained for subsequent analysis.

Mean normalized expression of the selected genes was calculated for homeostatic microglia and each ordered *Gpnmb* expression bin using Seurat AverageExpression(). Expression values were z-score scaled separately for each gene across the ordered groups. Genes were hierarchically clustered using a correlation-based distance matrix (1 – Pearson correlation) and Ward’s method (ward.D2). The resulting dendrogram was divided into four gene clusters (P1–P4), representing distinct expression programs across homeostatic microglia and progressively increasing *Gpnmb* expression within cluster Microglia_3.

For visualization of the average expression pattern of each gene cluster, z-score-scaled expression values were summarized across genes within P1–P4 for each ordered group. Lines represent the mean expression z-score of genes within each program, with the interquartile range shown as a ribbon.

Functional enrichment analysis was performed separately for genes assigned to P1–P4 using clusterProfiler::enricher and a custom TERM2GENE collection combining Gene Ontology Biological Process, Cellular Component, and Molecular Function terms with Hallmark gene sets. The background gene universe comprised genes detected in the analyzed microglial cells with total counts >20. Multiple-testing correction was performed using the Bonferroni method, and terms with adjusted *p* < 0.05 were considered significantly enriched. Enriched terms were ranked primarily by adjusted *p*-value and visualized according to gene count, gene ratio, and adjusted *p*-value.

### Transcription factor binding motif analysis

To infer candidate transcriptional regulators of microglia cluster–specific gene programs, cluster-specific marker genes were ranked by average log2 fold-change. For each cluster, the top 300 marker genes were selected and clusters annotated as microglia were retained. Motif enrichment analysis was performed for each microglia cluster using RcisTarget (v1.23.1), applying the 500 bp motif ranking database and restricting transcription factor (TF) annotations to high-confidence direct motif. Enrichment results were filtered to remove low-confidence TF assignments and summarized by retaining the motif with the highest normalized enrichment score (NES) per TF. The top enriched TFs (by NES) were visualized per microglia cluster in a dot plot (point size = number of enriched genes; color = NES). For each microglia cluster, TF regulatory networks were constructed by generating an edge list linking enriched motifs/TFs to their predicted target genes. Networks were visualized using igraph and ggraph, with force-directed layouts. Motif/TF nodes were annotated by the number of predicted targets and labeled with the corresponding TF name, while target gene nodes were displayed separately.

### Signature enrichment analysis

To quantify microglial activation and phenotype programs, published microglia gene signatures were compiled from multiple sources (Suppl. Table 2 & 3): (i) ^7^ disease-associated microglia (DAM) and homeostatic (HOMEO) genes, derived from the authors’ differential expression table by filtering for FDR < 0.05 and then sorting by absolute fold-change and FDR to extract the top 50 genes for both statistics respectively; (ii) ^8^ DAM and disease/inflammatory microglia (DIM) gene sets; (iii) ^9^ homeostatic microglia signature; and (iv) ^10^ signatures including HOMEO and DAM. For downstream scoring, signatures were merged into unified HOMEO, DIM, and DAM gene lists and filtered to retain genes expressed in at least 25% of cells in the dataset. Signature activity per cell was quantified using Seurat’s AddModuleScore, generating module scores for each state (HOMEO, DIM, DAM).

### Re-analyses of published data set: Depp et al. ^11^

Raw gene-level count data for microglia isolated from mouse brain in the Cnp-null and 5xFAD backgrounds (GSE178296) were downloaded and imported into R (v4.3.2). Counts were analyzed using DESeq2 (v1.42.1), including pre-filtering to retain genes with a total count sum ≥10 across all samples. Differential expression modeling and normalization were performed using the DESeq2 workflow, and normalized counts were used for downstream signature-based analyses.

Gene Set Variation Analysis (GSVA) was performed on the normalized expression matrix, filtered to retain genes with a row sum >50 across all samples and collapsed to unique gene symbols, to calculate per-sample enrichment scores for the P3 gene program defined in Figure 4 restricted to genes present in the dataset. GSVA scores were compared across genotypes using boxplots with overlaid jittered sample points. Group differences were assessed using pairwise two-sided Student’s *t*-tests.

### Re-analyses of published data set: Lengerich et al. ^12^

Publicly available single-cell RNA-seq data from Lengerich et al. (GSE209912) were downloaded from GEO and reprocessed in R using Seurat (v5.0.3). The raw sparse count matrix and associated barcode, gene symbol, and cell-level metadata files were imported and used to generate a Seurat object (min.features = 200, min.cells = 5). Gene expression values were log-normalized (LogNormalize, scale factor 10,000).

To quantify the transcriptional programs defined in the GPNMB-ordered analysis within the Lengerich et al. dataset, the P1–P3 gene sets identified in Figure 4 were used as independent signatures. Per-cell module scores for each signature were calculated using Seurat’s AddModuleScore() function. The resulting P1, P2, and P3 module scores were compared across experimental groups and visualized using violin plots.

### Re-analyses of published data set: Lee et al. ^13^

Publicly available single-cell RNA-seq data from Lee et al. (GSE153895) were downloaded from GEO and reprocessed in R using Seurat (v5.0.3). The raw sparse count matrices and associated barcode, gene symbol, and cell-level metadata files were imported per sample and used to generate a merged Seurat object (min.features = 200, min.cells = 5). Gene expression values were log-normalized (LogNormalize, scale factor 10,000), variable features were identified using the vst method, and data were scaled prior to dimensionality reduction. PCA was performed on variable genes and the first 20 principal components were used for UMAP embedding. Subsequently, cells were annotated through label transfer using published murine brain single-cell data sets^5^ and only cells identified as microglia were retained for further analyses. The microglia subset was re-normalized, variable genes were re-identified, scaled, and re-embedded (PCA/UMAP) followed by graph-based clustering (resolution 0.4). Cluster-specific marker genes were identified using Seurat’s FindAllMarkers (Wilcoxon rank-sum test; min.pct = 0.2, logfc.threshold = 0.2, min.diff.pct = 0.1, positive markers only). A cluster enriched for T cell-associated features was removed prior to downstream signature scoring. To quantify the transcriptional programs defined in the GPNMB-ordered analysis within the Lee et al. dataset, the P1–P3 gene sets identified in Figure 4 were applied as independent signatures. Per-cell enrichment of each signature was calculated using Seurat’s AddModuleScore() function, yielding P1, P2, and P3 module scores for each microglial cell. Module scores were compared across the indicated genotypes and visualized using violin plots.

### Re-analyses of published data set: Reich et al. ^14^

Raw bulk RNA-seq gene-level count data and sample annotations were obtained from GEO (GSE235515) and imported into R (v4.3.2). Lowly expressed genes were filtered out by retaining only genes with a total count sum ≥10 across all samples. A DESeq2 dataset was generated from the filtered count matrix with condition as the main factor, and the standard DESeq2 workflow was used for library size normalization, dispersion estimation, and negative binomial model fitting. Differential expression testing was performed for predefined pairwise contrasts. Multiple testing correction was performed using IHW, and effect sizes were optionally shrunk (normal prior). To assess the transcriptional programs defined in Figure 4 in the bulk RNA-seq dataset from Reich et al., the P1–P4 gene sets derived from the GPNMB-ordered microglial analysis were restricted to genes present in the dataset and used for GSVA. GSVA enrichment scores were calculated from DESeq2-normalized counts for each sample. Enrichment scores were compared across *WT* control, *WT* AAV-PGRN, *DKO* control, and *DKO* AAV-PGRN groups using boxplots with overlaid jittered sample points. Group differences were assessed using pairwise two-sided Student’s *t*-tests.

### NanoString targeted gene expression profiling

Snap-frozen brain tissue was mechanically powdered in liquid nitrogen. Qiagen RNeasy Mini Kit was employed for total RNA isolation. 100 ng RNA per sample were utilized for gene expression analysis with the NanoString Glial Profiling panel. Raw nCounter counts were imported into R (v4.3.2) using NanoStringNCTools (v1.10.0) and processed using NanoStringQCPro functions (v1.30.0) for initial quality control and normalization.

To further account for technical variability and unwanted sources of variation across samples, raw count matrices were subjected to RUV-based normalization using RUV_total() as implemented in the NanoNormIter package (v0.1.0). Briefly, genes were annotated according to NanoString probe classes and a factor of unwanted variation was estimated (k = 1). The resulting adjusted expression matrix was then used for downstream differential expression analysis. Differential expression was performed using DESeq2 (v1.42.1). A DESeqDataSet was generated from the RUV-normalized count matrix, restricting the analysis to endogenous genes only, and phenotype metadata were incorporated as sample-level covariates. Differential expression was assessed using a negative binomial generalized linear model in which gene expression was modeled as a function of genotype. DESeq2 was run using the standard workflow, including estimation of size factors, dispersion estimation, and Wald testing for genotype-associated expression differences.

### GSVA-based enrichment of microglia marker programs

To relate the transcriptional programs defined in the single-cell analysis to bulk NanoString expression profiles, the P1–P3 gene sets identified in the GPNMB-ordered analysis in Figure 4 were restricted to genes represented on the NanoString panel. GSVA was then performed on the normalized expression matrix to calculate per-sample enrichment scores for each signature. Enrichment scores were compared across the indicated genotypes and disease models using boxplots with overlaid individual samples. Group differences were assessed using pairwise two-sided Student’s *t*-tests.

For the relationship between the GPNMB-associated signature and neurodegeneration, mean P3 GSVA enrichment scores were calculated for each experimental group and correlated with mean plasma NfL levels using Pearson correlation. Linear regression was used to visualize the relationship, and mean ± SEM values were displayed for both P3 enrichment and NfL.

### AAV-PGRN rescue

Snap frozen spinal cord material from the Reich et al. 2024 study were used for protein extraction and analyzed by immunoblotting. In brief *DKO* mice received AAV-PGRN or control AAV via tail-vein injection at 6–8 weeks of age. Mice were analyzed at 3-4 months of age by immunoblotting to assess rescue of lysosomal dysfunction and modulation of DAM markers.

### Isolation of brain nuclei for ChIP-sequencing

Brain nuclei from *DKO* and *WT* mice were isolated as previously described ^15,16^, with initial homogenization performed with 1% formaldehyde in PBS for 10 min, followed by 0.125M glycine quench to halt the formaldehyde fixation. Nuclei were stained overnight at 4°C with PU.1-PE (Cell Signaling 81886S) and NEUN-AF488 (Millipore MAB 377X). Nuclei were washed with FACS buffer (10 mM Tris-HCl pH 8.0, 1 mM EDTA, 5 mM MgCl2, 0.1 M sucrose and 0.5% Triton X-100), passed through a 40 μm strainer, and stained with 0.5 μg/mL DAPI (BioLegend 422801). PU.1^+^ nuclei were sorted with a Beckman Coulter MoFlo Astrios EQ and pelleted at 1600xg for 5 min at 4°C in FACS buffer. Nuclei pellets were snap frozen and stored at -80°C prior to library preparation.

### H3K27ac ChIP-seq

Chromatin immunoprecipitation sequencing was performed as previously described ^17,18^. *For H3K27ac ChIP-seq*, ∼200000 fixed nuclei were quickly thawed on ice and resuspended in ice-cold LB3 buffer (10 mM Tris-HCl pH 7.5, 100 mM NaCl, 1 mM EDTA, 0.5 mM EGTA, 0.1% Na-Deoxycholate, 0.5% N-lauroylsarcosine) and 1x protease inhibitor cocktail. Chromatin was sheared by sonication using the PIXUL Multi-Sample Sonicator (Active Motif) in low bind 96 well plates (Corning, #7007) following the manufacturer’s instructions. Samples were sonicated with the following settings: Pulse (N): 50, PRF [kHz]: 1.00, Process time: 60 min, Burst Rate [Hz]: 20.00. Samples were pulse spun to 600xg at 4°C in the 96-well plate, transferred to low-bind tubes, and chromatin was recovered by retaining the supernatant following a spin at 21000xg at 4°C for 10 min. The supernatant was then diluted 1.1-fold with ice-cold 10% Triton X-100. Two percent of the lysate was kept as ChIP input control. 10 μL of Dynabead Protein A and 10 μL of Protein G were added per sample, in addition to 2 μg of a specific antibody for H3K27ac (Active Motif 39685). The samples were rotated overnight at 4°C and were washed as follows the next day: 3x with Wash Buffer I (20 mM Tris-HCl pH 7.5, 150 mM NaCl, 2 mM EDTA, 0.1% SDS, 1% Triton X-100) + 1x protease inhibitor cocktail, 3x with Wash Buffer III (10 mM Tris-HCl pH 7.5, 250 mM LiCl, 1% Triton X-100, 1 mM EDTA, 0.7% Sodium Deoxycholate) + 1x protease inhibitor cocktail, 2x with TET (0.2% Tween-20/TE) + 1/3x protease inhibitor cocktail, and 1x with IDTET (0.2% Tween-20, 10 mM Tris pH 8, 0.1 mM EDTA). Samples were finally resuspended in 25 μL TT buffer (10 mM Tris pH 8, 0.05% Tween 20) prior to on-bead library preparation.

Libraries were prepared with NEBNext Ultra II DNA library prep kit (NEB, E7645) reagents according to the manufacturer’s protocol on beads suspended in 25 μL TT (10 mM Tris/HCl pH 7.5, 0.05% Tween-20), with reagent volumes reduced by half. DNA was eluted and crosslinks reversed by adding 4 μL 10% SDS, 4.5 μL 5 M NaCl, 3 μL EDTA, 4 μL EGTA, 1 μL proteinase K (20 mg/ml), 16 μL water, incubating for 1 h at 55°C, then 30 min to overnight at 65°C. DNA was purified using 2 μL of SpeedBeads (Cytiva GE45152105050250) in a 20% PEG8000 / 1.5M NaCl solution to final concentration of 12% PEG, then eluted with 25 μL TT. DNA contained in the eluate was then amplified for 12-14 cycles in 25 μL PCR reactions using NEBNext High-Fidelity Q5 2X PCR Master Mix (NEB) and 0.5 mM each of primers Solexa 1GA and Solexa 1GB. Resulting libraries were size selected by gel excision to 200-500 bp, purified, and single end sequenced using an Illumina NovaSeq instrument.

### H3K27ac ChIP-seq analysis

ChIP-seq data were mapped with Bowtie2 (v2.3.5.1) ^19^. After mapping, HOMER (v4.11) ^17^ was used to convert reads aligning to a single genomic location into “tag directories” for subsequent analysis using the command makeTagDirectory. To quantify differential regions of H3K27 acetylation between genotypes, previously published wild type microglia ATAC-seq peaks ^20^ were merged with HOMER’s mergePeaks command. Merged ATAC-seq peaks were then annotated with raw H3K27ac tag counts from each sample with HOMER’s ^17^ annotatePeaks using parameters -noadj and -size 1000. Subsequently, DESeq2 was used to identify the differential H3K27ac signal with the cutoffs of FC>2 and p.adj<0.05 using getDiffExpression.pl. To identify motifs enriched in differential peak regions over background, HOMER’s motif analysis (findMotifsGenome.pl) using *de novo* motifs and the following parameters (-size 200 -len 8,10,12 -mis 3 -p 8) was used. Background sequences were comprised of random GC-matched genomic sequences. HOMER was also used to generate sequencing-depth normalized bedGraph/bigWig files for visualization in the UCSC Genome Browser (http://genome.ucsc.edu).

### Scanning electron microscopy (SEM)

Brains were perfusion fixed (4% PFA and 2.5% glutaraldehyde in 0.1 M sodium cacodylate buffer, pH 7.4; Science Services) and vibratome sections immersion fixed for 24h. We applied a standard rOTO en bloc staining protocol ^21^ including postfixation in 2% osmium tetroxide (EMS), 1.5% potassium ferricyanide (Sigma) in 0.1 M sodium cacodylate (Science Services) buffer (pH 7.4). The staining was enhanced by reaction with 1% thiocarbohydrazide (Sigma) for 45 min at 40°C. The sections were washed in water and incubated in 2% aqueous osmium tetroxide, washed and then further contrasted by overnight incubation in 1% aqueous uranyl acetate at 4°C and 2h at 50°C. Samples were dehydrated in an ascending ethanol series and infiltrated with LX112 (LADD). Blocks were cured for 48 hours, trimmed (TRIM2, Leica) and sectioned at 100 nm thickness using a 35° ultra-diamond knife (Diatome) on an ultramicrotome (UC7, Leica). Sections were collected onto ∼1 x 0.5 cm carbon coated Kapton tape strips (kindly provided by Richard Schalek and Jeff Lichtman, Harvard). The samples on tape were attached to adhesive carbon tape (Science Services) on 4-inch silicon wafers (Siegert Wafer) and grounded by adhesive carbon tape strips (Science Services). Scanning EM (SEM) micrographs were acquired on a Crossbeam Gemini 340 SEM (Zeiss) with a four-quadrant backscatter detector at 8 kV using ATLAS5 Array Tomography (Fibics). Medium lateral resolution images allowed the identification of regions of interest that were in turn imaged at 4-10 nm lateral resolution. Image analysis was performed in ImageJ ^22^.

### CSF collection and proteomics

Cerebrospinal fluid (CSF) was collected by cisterna magna puncture on deep anesthetized mice ^23^. Samples with visible blood contamination were excluded. Mouse CSF proteomics have been performed according ^23^. After CSF collection, mice were perfused and used for immunohistochemistry.

5 µL murine CSF was mixed with 2.75 µL denaturation buffer (0.5% Sodium Deoxycholate (5 mg/mL, SDC) in 250 mM ammonium bi-carbonate (19.76 mg/mL; NH4HCO3) and proteins were reduced and alkylated by with 2 µL 10 mM dithiothreitol (DTT) in 50 mM (ABC) for 30 min and 2 µL 55 mM iodoacetamide (IAA) for 20 min. After quenching with additional DTT, samples were digested with 0.1 µg LysC and 0.1 µg trypsin in 0.1% SDC and 50 mM ABC for 16 hrs at 37°C. Digestion was terminated by acidification with 5 % final formic acid (FA) and SDC removed by centrifugation at 16,000 x g for 10 min at 4 °C. Supernatants were desalted by STAGE Tip purification as previously published (Rappsilber et al., 2007). In brief, samples were loaded on two pre-equilibrated and washed C18 Empore disks, washed 3 times with 0.1% FA and eluted with 40 µL of 80% ACN with 0.1% FA. Samples were concentrated with vacuum centrifugation, reconstituted with 0.1% FA and 1/4 of the peptide mix was separated on a nanoElute system (Bruker, Germany) coupled to a TimsTOF pro mass spectrometer (Bruker, Germany) with a CaptiveSpray ion source (Bruker, Germany). Separation was performed on a self-packed C18 analytical column (15 cm (*DKO* mice) or 30 cm (*Npc1^-/-^* mice) × 75 µm ID, ReproSil-Pur 120 C18-AQ, 1.9 µm, Dr. Maisch GmbH, Germany) using a binary gradient of water (A) and acetonitrile (B) containing 0.1% FA at a flow rate of 250 nL/min (*Npc1^-/-^* mice) or 300 nL/min (*DKO* mice) and a column temperature of 50°C (0 min. 2% B, 3.5 min 5% B, 38 min 24% B, 59 min 35% B, 64 min 60% B). Mass spectra were acquired using data independent acquisition parallel accumulation–serial fragmentation (DIA-PASEF) in combination with ion accumulation and separation using trapped ion mobility spectrometry (TIMS). For *Npc1^-/-^*CSF, one MS1 full scan was followed by 2 rows of 42 sequential DIA windows with 24 m/z width for peptide fragment ion spectra with an overlap of 2 m/z covering a scan range of 350 to 1275 m/z. The ramp time was fixed to 166 ms and 6 windows were scanned per ramp. This resulted in a total cycle time of 2.5 s. For *DKO* mice, a ramp time of 100 ms was set. One scan cycle included one MS1 full scan, followed by 26 windows with a width of 27 m/z covering a m/z range of 350–1001 m/z. Two windows were recorded per PASEF scan at a cycle time of 1.4 s in positive mode.

Samples were analyzed with DIA-NN (version 1.9.2) ^24^ using a library-free search against a full mouse proteome (derived from Uniprot). Number of missed cleavages was set to 2, peptide length to 7 – 30 and precursor charges to 2 – 4. Cysteine carbamidomethylation was set as a fixed modification, the number of variable modifications to 2 and included UniMod:35 and UniMod:1. Match-between-runs was enabled, cross-run normalization was disabled. Statistical analysis was performed using Perseus and Python. In short, after filtering for protein groups present with minimum 2 peptides in 3 out of 4 replicates and removal of typical contaminants, missing values were replaced by downshift imputation.

### Analysis of *DKO* and *Npc1^-/-^* proteomics overlap and functional CSF enrichment

To identify shared proteomic alterations between the *DKO* and *Npc1*^-/-^, the list of overlapping differentially abundant proteins (Suppl. Table 6, 7, 8, 9) was extracted and Gene Ontology enrichment analysis was performed using the clusterProfiler package. GO term-to-gene mappings for biological process (GO:BP), cellular component (GO:CC), and molecular function (GO:MF) were combined into a single annotation table and over-representation analysis was conducted using enricher() with the overlapping protein list as input and the combined GO reference annotation. Multiple testing correction was applied using the Bonferroni method. GO terms were considered significantly enriched at adjusted p-value and q-value thresholds of 0.05. Enrichment results were visualized using the emapplot.

### Quantification and statistical analysis

Statistical analyses were performed using GraphPad Prism 10, R v4.3.2 and v4.3.3 as indicated, and Python. Data are presented as mean ± SEM unless noted. For comparisons between two groups, unpaired two-tailed Student’s t-test were used if not otherwise stated. For multi-group comparisons, one-way ANOVA with Tukey’s post hoc test was applied. Significance thresholds were set at p < 0.05 unless otherwise indicated. Exact statistical tests, n values, and p values are reported in figure legends. Plots were generated using ggplot2 if not otherwise indicated.

### CLEC7A antibody generation

For generation of monoclonal antibodies against mouse CLEC7A, a Lou/c rat was immunized subcutaneously (s.c.) and intraperitoneally (i.p.) with a mixture of 40 µg ovalbumin-coupled peptide corresponding to mClec7a aa 16-36 TQLDFSTQDIHKRPRGSEKGS (Peps4LS, Heidelberg) dissolved in 400 µl PBS containing 5 nmol CpG2003 (TIB MOLBIOL, Berlin, Germany), and 400 µl incomplete Freund’s adjuvant (Sigma-Aldrich). After 12 weeks, a boost without Freund’s adjuvant was given i.p. and s.c. 3 days before fusion. Fusion of mouse immune spleen cells with the myeloma cell line P3X63-Ag8.653 was performed using polyethylene glycol 1500 according to standard procedure (Koehler and Milstein, Nature. 1975, 256:495-497). After fusion, the cells were plated in 96-well plates in RPMI 1640 medium supplemented with 20% fetal calf serum, 1mM pyruvate, 1x non-essential amino acids and HAT media supplement (Hybri-Max, Sigma-Aldrich).

Hybridoma supernatants were screened 10 days later in a flow cytometry assay (iQue, Intellicyt; Sartorius) using biotinylated peptides captured on streptavidin beads (PolyAN, Berlin) and incubated for 90 min with hybridoma supernatant and Atto-488-coupled isotype-specific monoclonal mouse-anti-rat IgG secondary antibodies. Antibody binding was analyzed using ForeCyt software (Sartorius). Positive supernatants were further validated by capture ELISA and immunoblotting. Hybridoma cells from selected supernatants were subcloned by limiting dilution to obtain stable monoclonal cells lines. Experiments in this work were performed with clone CLCMA 1F1 (ratIgG2a/k). Animal procedures were conducted in accordance with the German Animal Welfare Law and with approval by the local authorities of Upper Bavaria, Germany (animal license: ROB-55.2Vet-2532.Vet_03-17-68).

### *Clec7a* KO mouse generation

Strategy to generate *Clec7a*^KO^ by CRISPR/Cas9-assisted gene editing in mouse zygotes as described previously ^25^. A CRISPR/Cas9-mediated Indel mutation was introduced in exon 1 of *Clec7a* leading to a frame-shift and early stop codon. *Clec7a*^KO^ RNP consisted of 640 ng/µl SpCas9 protein (IDT), 4 µM crClec7aEx1 (IDT), and 4 µM tracrRNA (IDT), and was electroporated into zygotes using the NEPA21 electroporator and a 1 mm electrode. After electroporation, treated zygotes were transferred to pseudopregnant foster animals. Pronuclear stage zygotes were obtained by mating C57BL/6J males with superovulated C57BL/6J females (Charles River). Embryos were then microinjected into the male pronucleus with an injection mix containing *Clec7a*-specific Cas9 ribonucleoproteins (RNP). After microinjection, treated zygotes were transferred to pseudo-pregnant foster animals. To exclude additional unwanted modifications, putative off-target sites were predicted using the online tool CRISPOR (http://crispor.tefor.net/) ^26^. For analysis, genomic DNA of WT and heterozygous *Clec7a*^KO/+^ mice was isolated and predicted loci with a CFD score > 0.40 and an MIT score > 0.7 were PCR-amplified and Sanger sequenced. Animals without any off-target mutations were used for generation of homozygous *Clec7a*^KO/KO^ strain. Animals were raised and handled at the DZNE Munich. All animal experiments were approved by the Bavarian government under license number ROB-55.2-2532.Vet_02-16-121. Oligonucleotides used for generation and verification of the Clec7a KO mouse line are listed in the following resource table.

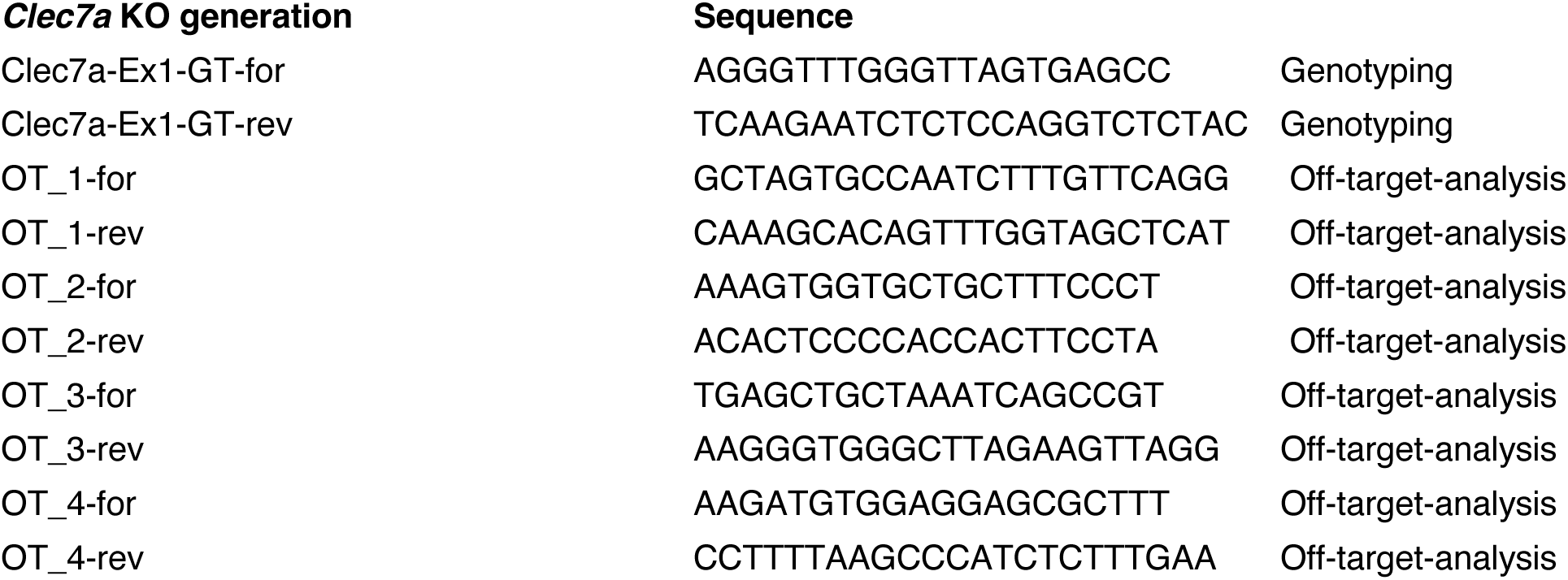

## Resource Table

| **REAGENT** | **SOURCE** | **IDENTIFIER** |
|---|---|---|
| ABI3 | Cell Signaling | CST 23060S |
| APOE AB947 | Chemicon® | AB947 |
| BAX | THERMO | MA5-35342 |
| CLEC7A / Dectin-1 | InvivoGen | R1-8g7 |
| CLEC7A inhouse | Regina Feederle | CLCMA 1F1 |
| CSF-1R/M-CSF-R | Cell Signaling | 3152 |
| CTSB | R&D Systems | AF965 |
| CTSD | R&D Systems | AF1029 |
| CTSL | R&D Systems | AF1515 |
| FABP5 (D1A7T) | Cell Signaling | 39926T |
| GALECTIN3 | Biolegend | 125402 |
| GFAP | Thermo | PA5-18598 |
| GPNMB (E7U1Z) | Cell Signaling | 90205 |
| GPNMB/Osteoactivin | R&D Systems | AF2330-SP |
| IBA1 | Synaptic | 234 009 |
| IBA1 | GeneTex | GTX100042 |
| LC3 | Novusbio | NB100 |
| LC3 (D3U4C) | Cell Signaling | 12741S |
| Lysozyme C-1/2 (E2K5R) | Cell Signaling | 60487T |
| MBP | Biolegend | 808401 |
| P62 (SQSTM1) | MBL | PM045 |
| Serpin A3N | R&D Systems | AF4709 |
| SPP1/ Osteopontin | Proteintech | 22952-1-AP |
| Tau (E8B5J) | Cell Signaling | 54926S |
| TDP43 (P) | Proteintech | 80007-1-RR |
| TDP43(C-term) | Proteintech | 12892-2AP |
| TMEM106B (E7H7Z) | Cell Signaling | #93334 |
| TREM2 | Thermo | MA5_31267 |
| TREM2 E7P8J | Cell Signaling | 76765S |
| UBIQUITIN | Santa Cruz | SC 3017 P4D1 |
| CD11b APC-CY7 | BD Bioscience | 557657 |
| O4 Antibody, PE | Miltenyi Biotec | 130-117-507 |
| ACSA-2 Antibody, PE | Miltenyi Biotec | 130-123-284 |
| Neuron Isolation Kit, mouse | Miltenyi Biotec | 130-115-390 |
| Adult Brain Dissociation Kit, mouse and rat | Miltenyi Biotec | 130-107-677 |
| BD Rhapsody Cartridge Kit | Becton Dickinson | 633733 |
| BD Cartridge Reagent Kit | Becton Dickinson | 633731 |
| BD Rhapsody cDNA Kit | Becton Dickinson | 633773 |
| BD Rhapsody™ WTA Amplification Kit | Becton Dickinson | 633801 |
| High Sensitivity D5000 ScreenTape | Agilent | 5067-4626 |
| Qubit dsDNA HS Assay Kit | ThermoFisher | Q32851 |
| NovaSeq 6000 S4 Reagent Kit (200 cycles) | Illumina | 20028313 |
| **DEPOSITED DATA** |  |  |
| scRNA-seq raw data | This paper | GEO |
| Processed scRNA-seq data | This paper | GEO |
| Analysis code | This paper | Gitlab |
| **SOFTWARE AND ALGORITHMS** |  |  |
| Bcl2fastq2 | Illumina | V2.20 |
| STAR | <sup>27</sup> | v2.7.3a |
| Cutadapt | <sup>28</sup> | v1.16 |
| Dropseq-tools | https://github.com/broadinstitute/Drop-seq/ | v2.0.0 |
| R | https://www.cran.r-project.org | v4.3.2; v4.3.3 |
| Seurat | <sup>29</sup> | v5.0.3 |
| DoubletFinder | <sup>30</sup> | v2.0.4 |
| harmony | <sup>31</sup> | v1.2.0 |
| DESeq2 | <sup>32</sup> | v1.42.1 |
| GSVA | <sup>33</sup> | v1.50.2; v1.50.5 |
| clusterProfiler | <sup>34</sup> | v4.10.1 |
| RcisTarget | <sup>35</sup> | v1.23.1 |
| NanoStringNCTools | <sup>36</sup> | v1.10.0 |
| igraph | https://igraph.org | v2.0.3 |
| ggraph | https://ggraph.data-imaginist.com | v2.2.1 |

---

**Fig. S1:**
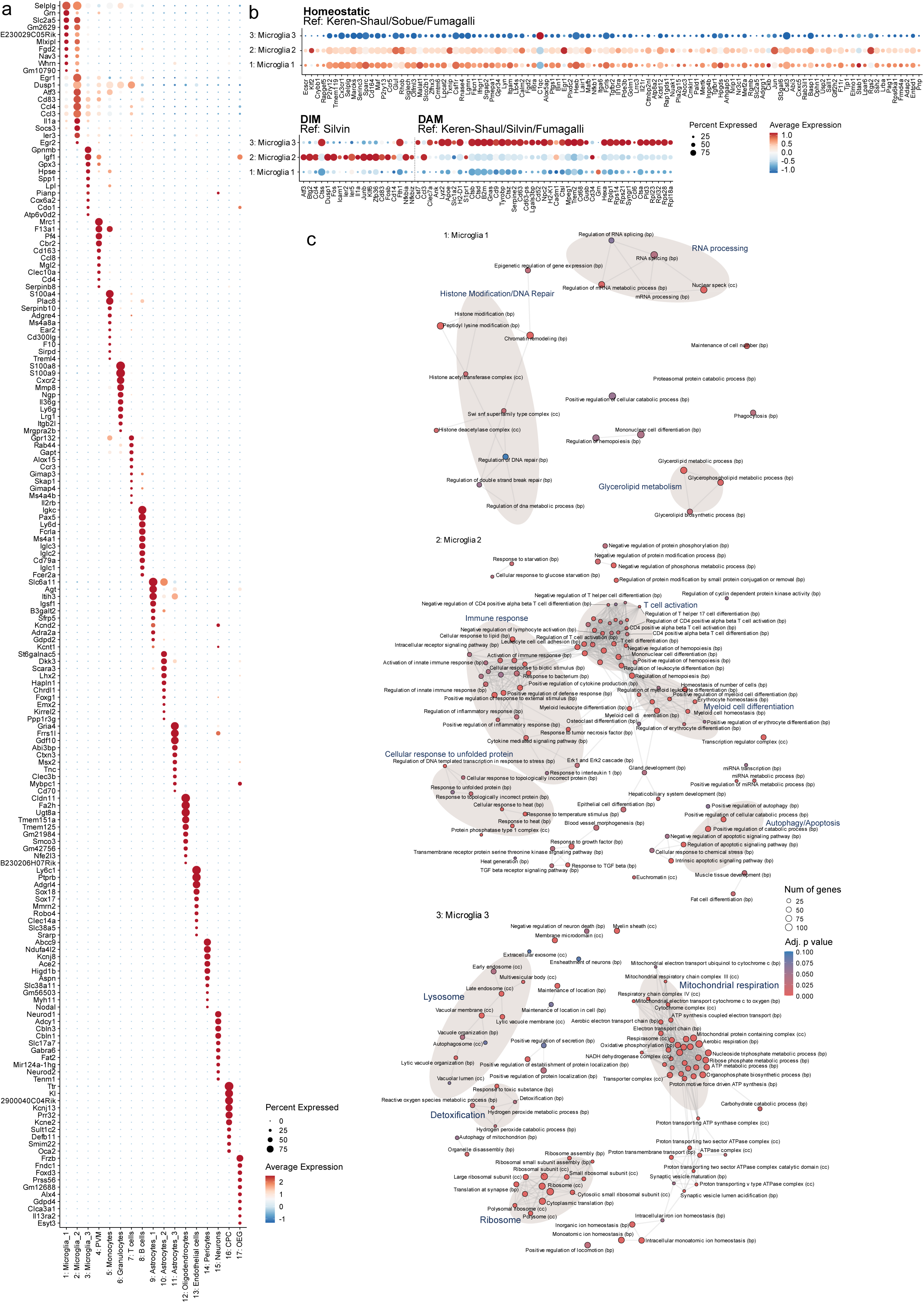
Cellular characterization of microglial phenotypes. (a) Dot plot of the top 10 marker genes per cluster sorted by average log fold change. (b) Dot plot of gene expression across microglia clusters 1, 2, and 3 for HOM, DIM and DAM microglia signatures derived from ^2,3,88,113^. (c) Network visualization of Gene Ontology enrichment results based on cluster marker genes (min.pct = 0.1, logfc.threshold = 0.2) of microglia cluster 1, 2 and 3 colored according to adjusted p-values and scaled according to the number of genes enriched per term. Edges indicate overlapping genes.

**Fig. S2:**
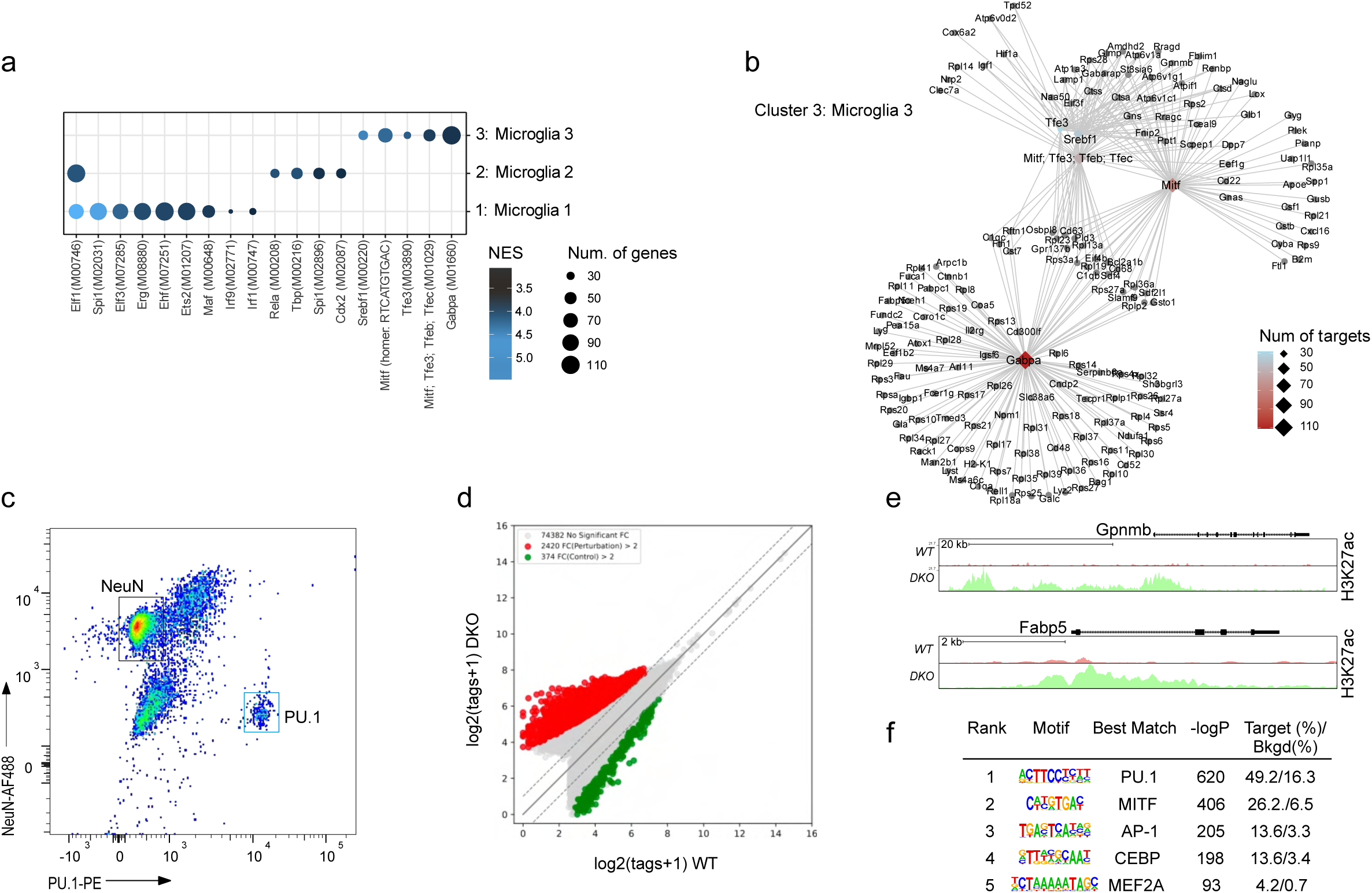
Enhancer remodeling in GPNMB⁺ DAM in *DKO* mice. (a) Dot plot showing transcription factor (TF) motif enrichment for cluster-specific marker genes (top 300 genes ranked by average log2 fold-change per cluster) identified using RcisTarget. Dot size indicates the number of predicted target genes associated with the TF motif, and color represents the normalized enrichment score (NES). (b) Network visualization of predicted transcriptional regulatory interactions based on enriched TF motifs from (A). Nodes represent transcription factors and their predicted target genes, and edges indicate TF–target relationships inferred from motif enrichment analysis. (c) Representative fluorescence-activated nuclei sorting (FANS) plot showing gating strategy used to collect PU.1^⁺^ nuclei for subsequent H3K27ac ChIP-seq and NeuN^⁺^ nuclei, which were excluded for downstream analysis. (d) Scatter plot depicting differential H3K27ac levels at open chromatin regions between *WT* and *DKO* PU.1^⁺^ microglia. Differential acetylation regions (DAR) with significantly increased H3K27ac in *DKO* are shown in red (fold change > 2, FDR <0.05, whereas regions with decreased H3K27ac are shown in green (fold change < -2, FDR < 0.05). (e) UCSC genome browser tracks showing H3K27ac ChIP-seq peaks at the top altered loci, *Gpnmb* (upper) and *Fabp5* (lower). (f) *De novo* motif enrichment analysis results of activated enhancers in *DKO* PU.1^⁺^ microglia.

**Fig. S3:**
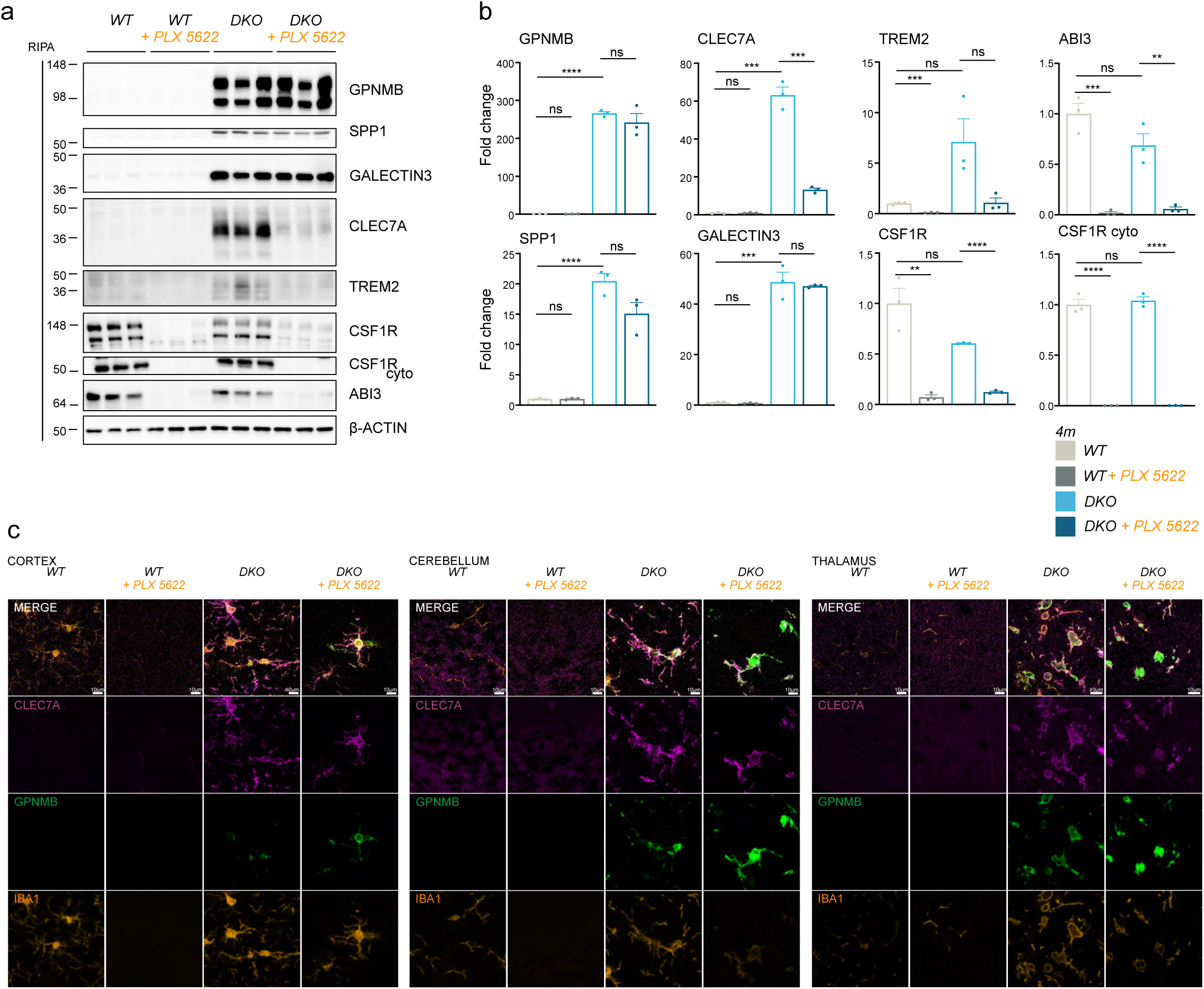
GPNMB⁺ DAM exhibit relative resistance to CSF1R-inhibition in *DKO* mice. (a) Western blot analysis of GPNMB, SPP1, GALECTIN3, CLEC7A, TREM2, CSF1R, ABI3 and β-ACTIN in the indicated fractions of total brain lysates from of 4-month-old *WT* and *DKO* mice ± PLX5622 treatment, (n = 3 mice per group). Asterisk indicates unspecific band. (b) Quantification of proteins in (D), fold change to *WT* (n = 3 mice per group). (c) Immunofluorescence of microglia activation markers CLEC7A(magenta), GPNMB(green) and IBA1(orange) in cortex, thalamus and the cerebellum of sagittal brain sections. Representative images of 4-month-old *WT* and *DKO* mice ± PLX5622 treatment. Scale bar indicates 10 µm.

**Fig. S4:**
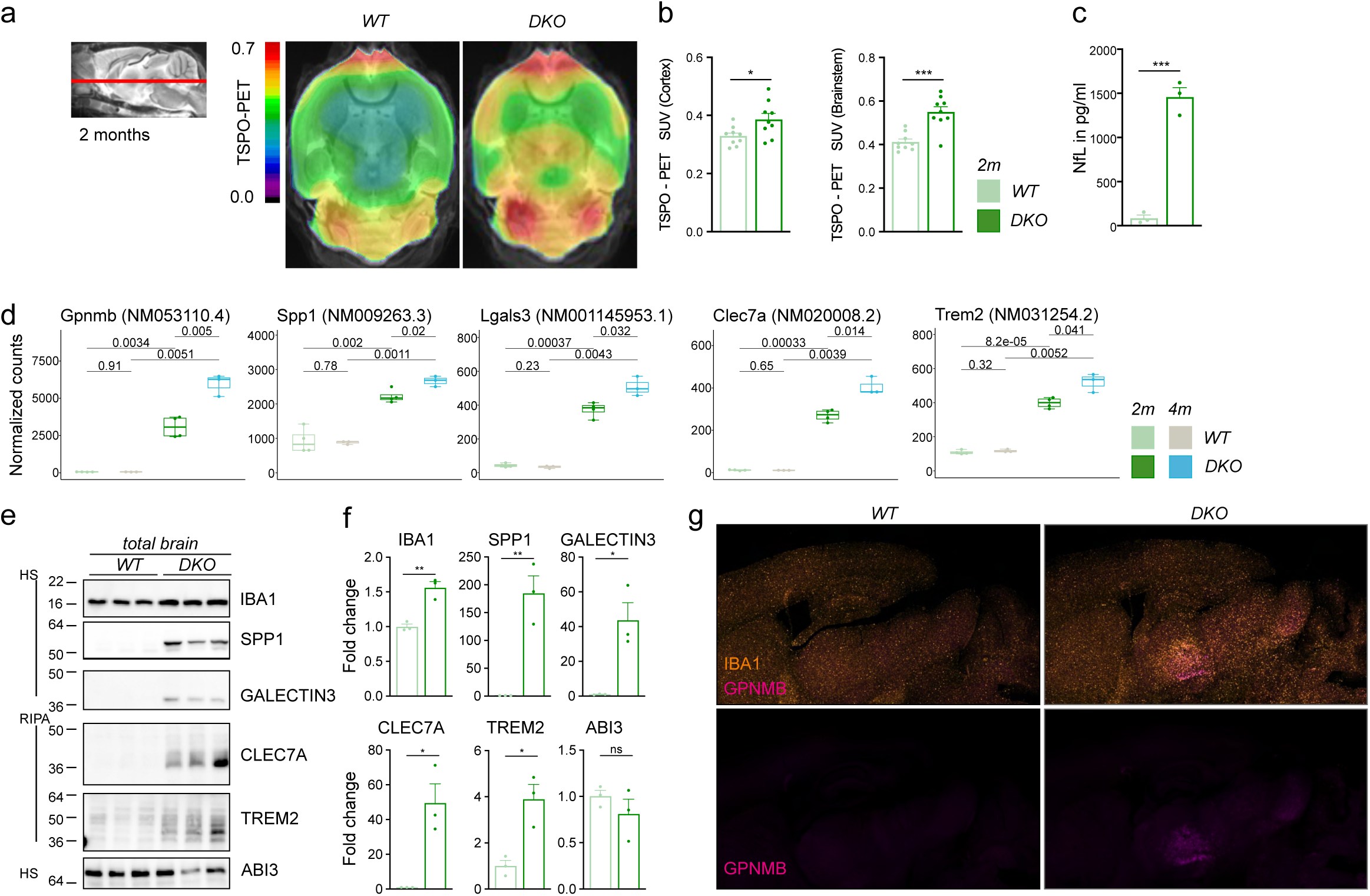
The GPNMB-associated program emerges during development of neurodegeneration. (a) *In vivo* ^18^F-GE180 TSPO µPET imaging. *WT* and *DKO* mice underwent TSPO µPET at 2 months of age. (b) Quantification of *in vivo* ^18^F-GE180 TSPO µPET imaging. Data represent the mean ± SD (n = 9 mice per group). (c) NfL in pg/ml of plasma of mice at 2 months (n = 3 mice per group). (d) Box plots of normalized counts of selected microglia genes in genes in total brain bulk transcriptomes of 2- and 4-month-old *WT* and *DKO* mice. P-values from two-sided Student’s *t*-tests are shown. (e) Western blot analysis of SPP1, GALECTIN3, CLEC7A, TREM2, and ABI3 in the indicated fractions of total brain lysates from of 2-month-old *WT* and *DKO* mice, (n = 3 mice per group). (f) Quantification of proteins in (e), fold change to *WT* (n = 3 mice per group). (g) Immunofluorescence of microglia activation markers GPNMB (magenta) and IBA1(orange), overview of sagittal brain sections. Representative images of 2-month-old *WT* and *DKO* mice treatment.

**Fig. S5:**
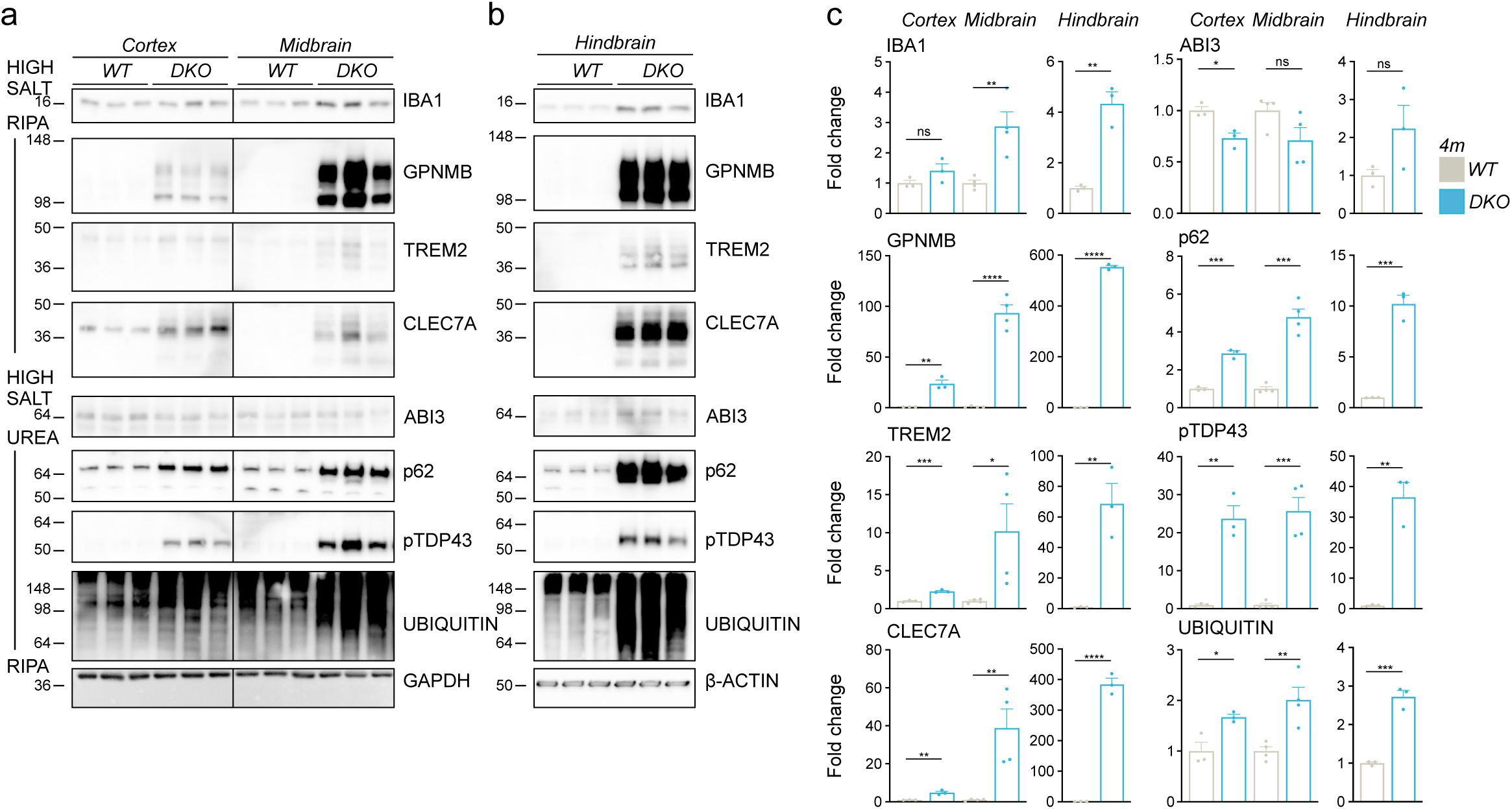
Regional GPNMB induction correlates with pathological protein accumulation. (a, b) Western blot analysis of GPNMB, TREM2, CLEC7A, ABI3, p62, pTDP43, UBIQUITIN and GAPDH or β-ACTIN in lysates from different brain regions of 4-month-old *WT* and *DKO* mice, (n = 3 mice per group). (C) cortex and midbrain, (d) hindbrain. (c) Quantification of proteins in (a, b), fold change to *WT* (n = 3 mice per group).

**Fig. S6:**
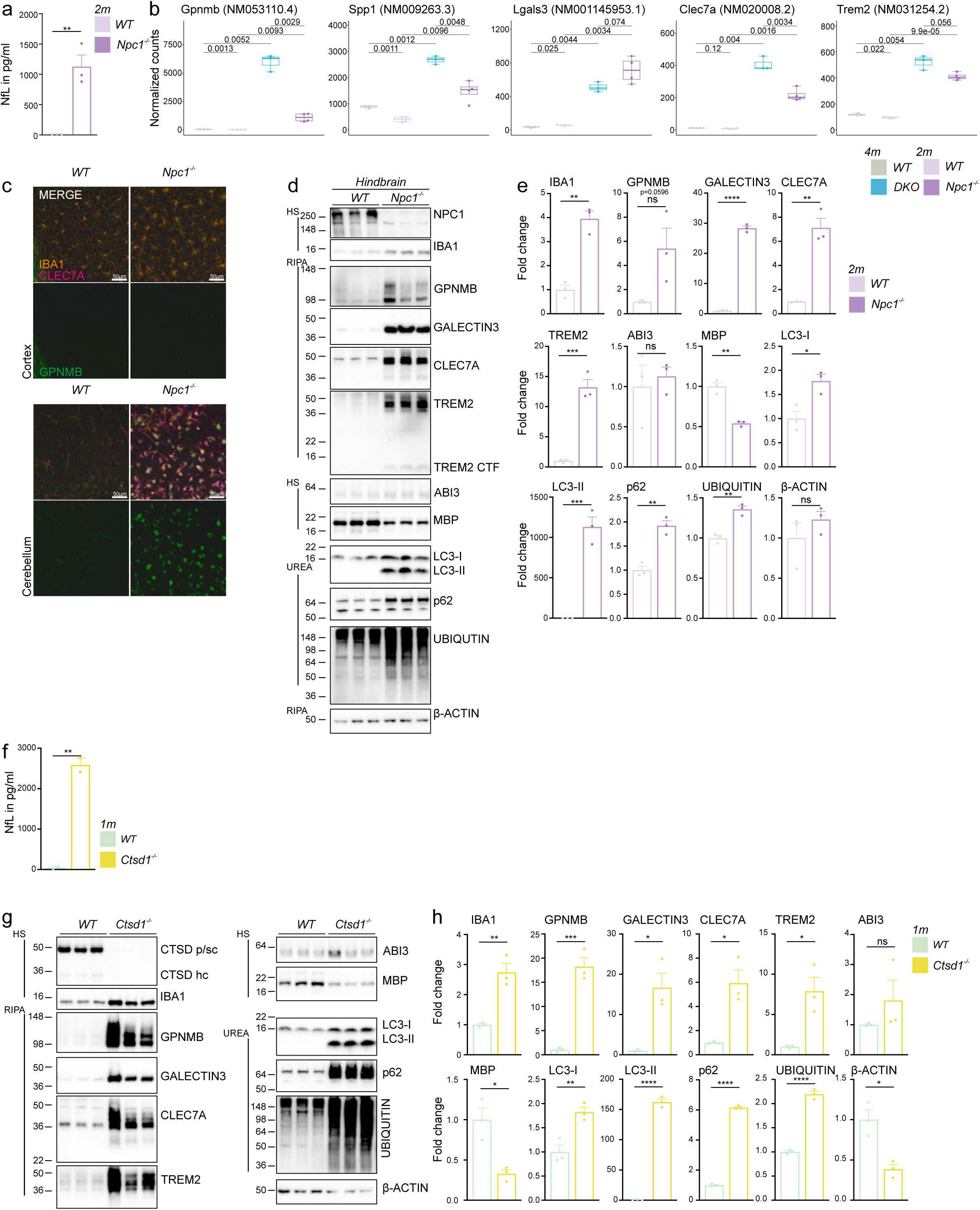
The GPNMB-associated program is consistently induced across mouse models with lysosomal dysfunction. (a) NfL in pg/ml of plasma of *WT* and *Npc1^-/-^* mice at 2 months (n = 3 mice per group). (b) Box plots of normalized counts of selected microglia genes in genes in total brain bulk transcriptomes of 2-month-old *WT* and *Npc1^-/-^* mice and 4-month-old *WT* and *DKO* mice. P-values from two-sided Student’s *t*-tests are shown. (c) Immunofluorescence of microglia activation markers CLEC7A (magenta), GPNMB (green) and IBA1 (orange) of sagittal brain sections. Representative images from cortex and cerebellum of 2-month-old *WT* and *Npc1*^-/-^ mice treatment. (d) Western blot analysis of NPC1, IBA1, GPNMB, GALECTIN3, CLEC7A, TREM2, ABI3, MBP, LC3, p62, UBIQUITIN and β-ACTIN in the indicated fractions of hindbrain lysates from *WT* and *Npc1^-/-^* mice at 2 months (n = 3 mice per group). (e) Quantification of proteins in (d), fold change to *WT* (n = 3 mice per group). (f) NfL in pg/ml of plasma of *WT* and *Ctsd^-/-^* mice at 1 month (n = 2 mice per group). (g) Western blot analysis of CTSD, IBA1, GPNMB, GALECTIN3, CLEC7A, TREM2, ABI3, MBP, LC3, p62, UBIQUITIN and β-ACTIN in the indicated fractions of total brain lysates from *WT* and *Ctsd^-/-^* mice at 1 month (n = 3 mice per group). (h) Quantification of proteins in (g), fold change to *WT* (n = 3 mice per group).

**Fig. S7:**
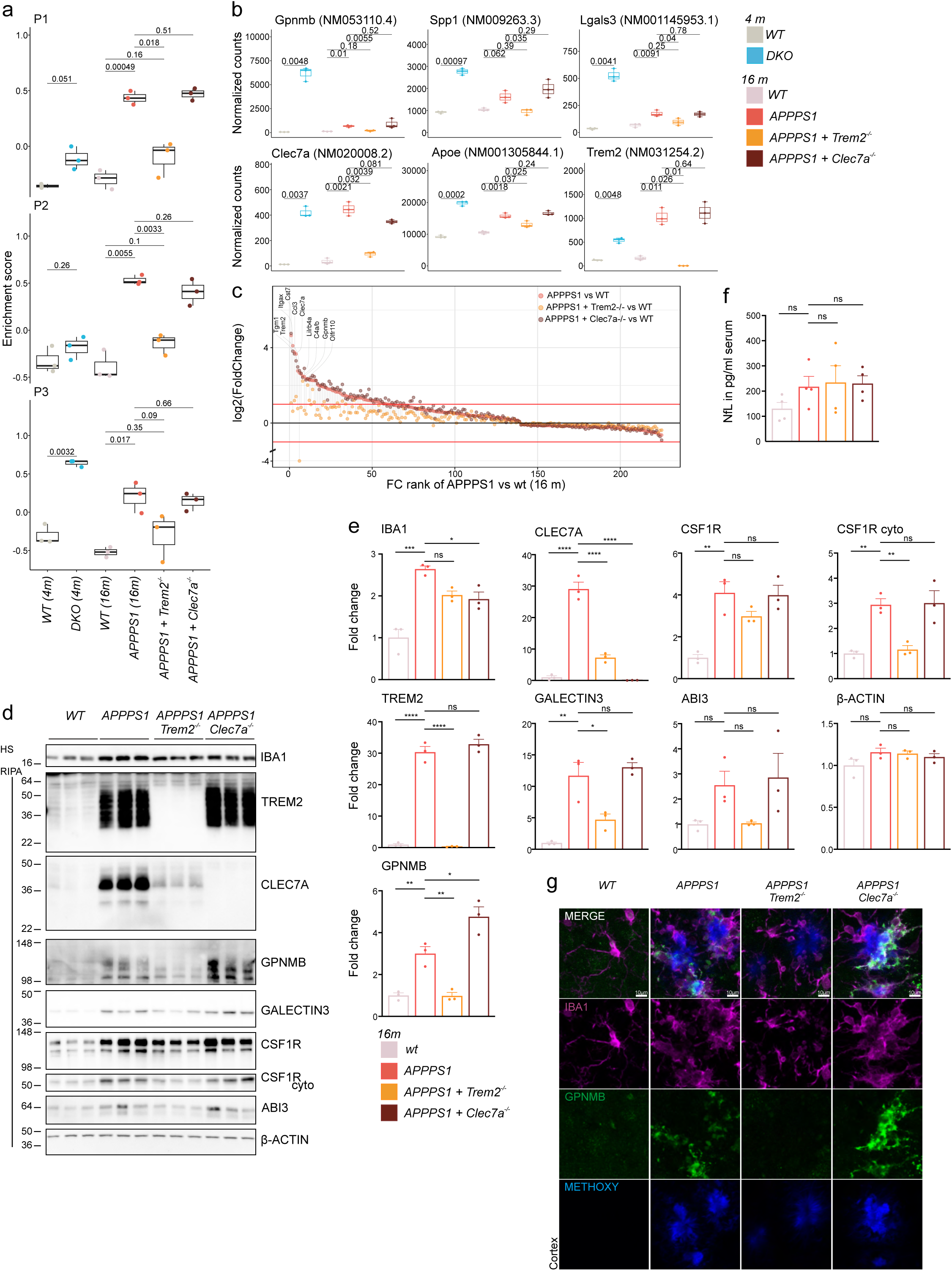
TREM2 regulates DAM activation and induction of the GPNMB-associated program in APPPS1 mice. (a) Box plots showing enrichment values of filtered microglia signatures for clusters 1, 2, and 3, in total brain bulk transcriptomes of 4-month-old DKO mice relative to 4-month-old WT controls and 16-month-old *APPPS1, APPPS1*x*Trem2^-/-^* and *APPPS1*x*Clec7a^-/-^*relative to 16-month-old *WT* controls. Microglia signature genes derived from scRNA-seq were filtered for minimal differential signal in bulk transcriptomics across the indicated comparisons (see methods). P-values from two-sided Student’s *t*-tests are shown. (b) Box plots of normalized counts of selected microglia genes in 4-month-old *WT* and *DKO* mice and 14-month-old *WT* and *APPPS1, APPPS1*x*Trem2^-/-^* and *APPPS1*x*Clec7a^-/-^* mice. P-values from two-sided Student’s *t*-tests are shown. *(c)* FC rank of 14-month-old *WT* and *APPPS1, APPPS1*x*Trem2^-/-^* and *APPPS1*x*Clec7a^-/-^* mice. (d) Western blot analysis of IBA1, TREM2, CLEC7A, GPNMB, GALECTIN3, CSF1R, ABI3 and β-ACTIN in the indicated fractions of total brain lysates from *WT, APPPS1, APPPS1*x*Trem2^-/-^* and *APPPS1*x*Clec7a^-/-^* . Mice at 14 months of age. (e) Quantification of proteins in (F), fold change to *WT* (n = 3 mice per group). (f) NfL in pg/ml of plasma of *WT, APPPS1, APPPS1*x*Trem2^-/-^* and *APPPS1*x*Clec7a^-/-^* mice at 14 months of age (n = 3 mice per group). (g) Immunofluorescence of microglia activation markers IBA1(magenta), GPNMB (green) and Ab-plaques by Methoxy-X04(blue) in the cortex of sagittal brain sections. Representative images of 14-month-old mice. Scale bar indicates 10 µm.

**Fig. S8:**
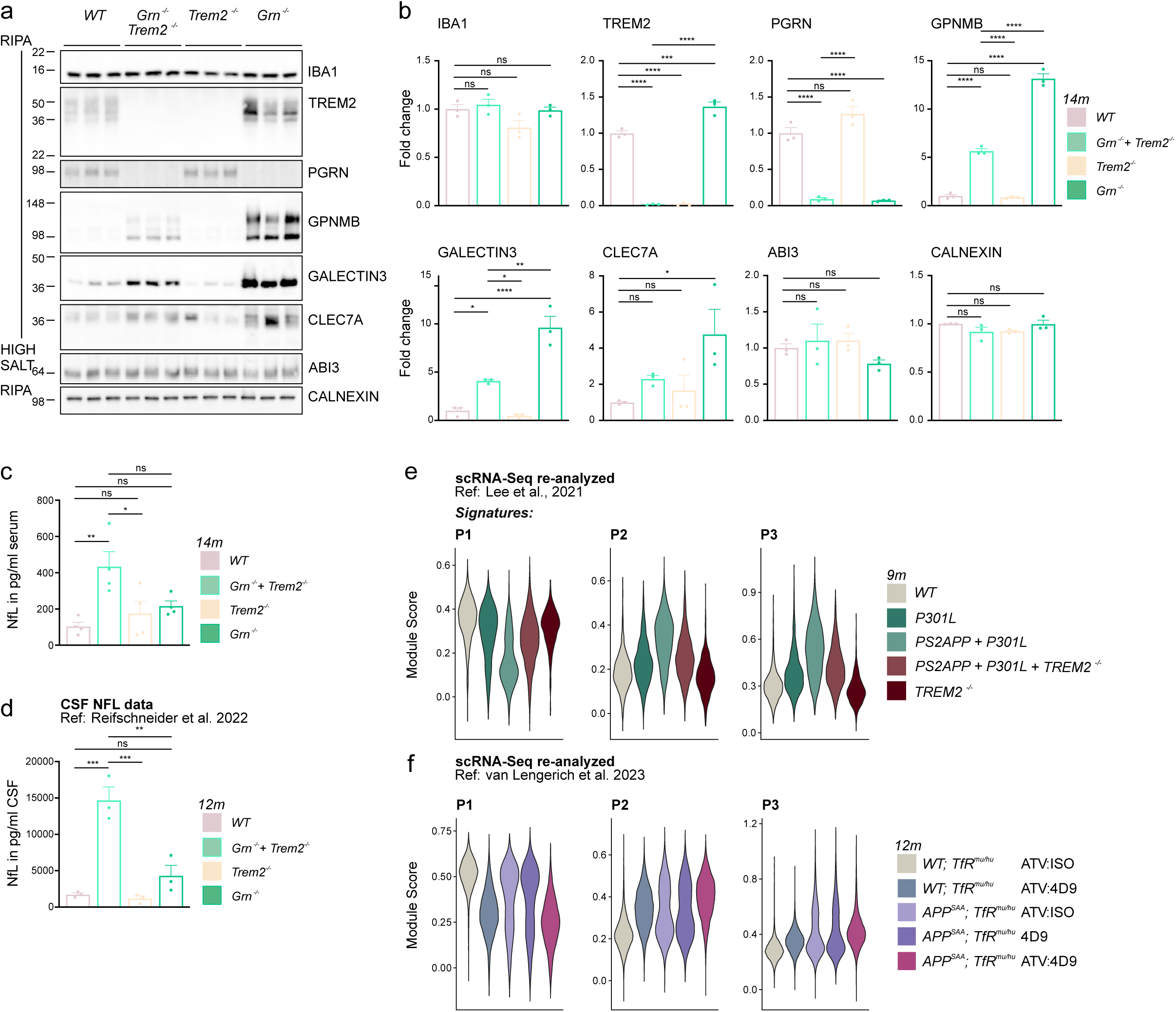
Loss of the TREM2 exacerbates neurodegeneration under lysosomal stress in independent disease models. (a) Western blot analysis of IBA1, TREM2, PGRN, GPNMB, GALECTIN3, CLEC7A, and ABI3 in the indicated fractions of total brain lysates from *WT, Grn^-/-^* x *Trem2^-/-^, Trem2^-/-^, Grn^-/-^* mice at 14 months of age (n = 3 mice per group). (b) Quantification of proteins in (D), fold change to *WT* (n = 3 mice per group). (c) NfL in pg/ml of plasma of *WT, Grn^-/-^* x *Trem2^-/-^, Trem2^-/-^, Grn^-/-^* mice at 14 months of age (n = 4 mice per group). (d) NfL in pg/ml of CSF of *WT, Grn^-/-^* x *Trem2^-/-^, Trem2^-/-^, Grn^-/-^* mice at 12 months of age (n = 3 mice per group), from ^78^. (e) Violin plots showing module scores of microglia cluster signatures based on scRNA-seq analysis shown in Fig.2 in microglia scRNA-seq data derived from *WT, P301L, PS2APPxP301L, PS2APPxP301LxTrem2^-/-^* mice at 9 months of age re-analyzed from ^79^. (f) Violin plots showing module scores of microglia cluster signatures based on scRNA-seq analysis from Fig. 2 in microglia scRNA-seq data derived from WT and APP^SAA^ mice expressing a transferrin receptor (TfR) binding site for the antibody transport vehicle (ATV) brainshuttle. Mice were treated with TREM2-stimulating ATV:4D9, 4D9, or ATV:ISO isotype control antibodies. Data were re-analyzed from ^24^.

## References

1. Butovsky, O. et al. Identification of a unique TGF-β-dependent molecular and functional signature in microglia. Nat. Neurosci. 17, 131–143 (2014).

2. Keren-Shaul, H. et al. A Unique Microglia Type Associated with Restricting Development of Alzheimer’s Disease. Cell 169, 1276–1290.e17 (2017).

3. Silvin, A. et al. Dual ontogeny of disease-associated microglia and disease inflammatory macrophages in aging and neurodegeneration. Immunity 55, 1448–1465.e6 (2022).

4. Park, M. D., Silvin, A., Ginhoux, F. & Merad, M. Macrophages in health and disease. Cell 185, 4259–4279 (2022).

5. Krasemann, S. et al. The TREM2-APOE pathway drives the transcriptional phenotype of dysfunctional microglia in neurodegenerative diseases. Immunity 47, 566–581.e9 (2017).

6. Hammond, T. R. et al. Single-Cell RNA Sequencing of Microglia throughout the Mouse Lifespan and in the Injured Brain Reveals Complex Cell-State Changes. Immunity 50, 253–271.e6 (2019).

7. Ulland, T. K. et al. TREM2 maintains microglial metabolic fitness in alzheimer’s disease. Cell 170, 649–663.e13 (2017).

8. Fracassi, A. et al. TREM2-induced activation of microglia contributes to synaptic integrity in cognitively intact aged individuals with Alzheimer’s neuropathology. Brain Pathol. 33, e13108 (2023).

9. Spangenberg, E. E. et al. Eliminating microglia in Alzheimer’s mice prevents neuronal loss without modulating amyloid-β pathology. Brain 139, 1265–1281 (2016).

10. Sosna, J. et al. Early long-term administration of the CSF1R inhibitor PLX3397 ablates microglia and reduces accumulation of intraneuronal amyloid, neuritic plaque deposition and pre-fibrillar oligomers in 5XFAD mouse model of Alzheimer’s disease. Mol. Neurodegener. 13, 11 (2018).

11. Lodder, C. et al. CSF1R inhibition rescues tau pathology and neurodegeneration in an A/T/N model with combined AD pathologies, while preserving plaque associated microglia. Acta Neuropathol. Commun. 9, 108 (2021).

12. Kiani Shabestari, S., et al. Absence of microglia promotes diverse pathologies and early lethality in Alzheimer’s disease mice. Cell Rep. 39, 110961 (2022).

13. Elmore, M. R. P. et al. Colony-stimulating factor 1 receptor signaling is necessary for microglia viability, unmasking a microglia progenitor cell in the adult brain. Neuron 82, 380–397 (2014).

14. Feiten, A. F. et al. TREM2 expression level is critical for microglial state, metabolic capacity and efficacy of TREM2 agonism. Nat. Commun. 17, (2026).

15. Schlepckow, K., Morenas-Rodríguez, E., Hong, S. & Haass, C. Stimulation of TREM2 with agonistic antibodies-an emerging therapeutic option for Alzheimer’s disease. Lancet Neurol. 22, 1048–1060 (2023).

16. Chen, Y. & Colonna, M. Microglia in Alzheimer’s disease at single-cell level. Are there common patterns in humans and mice? J. Exp. Med. 218, (2021).

17. Atagi, Y. et al. Apolipoprotein E is a ligand for triggering receptor expressed on myeloid cells 2 (TREM2). J. Biol. Chem. 290, 26043–26050 (2015).

18. Wang, Y. et al. TREM2 lipid sensing sustains the microglial response in an Alzheimer’s disease model. Cell 160, 1061–1071 (2015).

19. Kleinberger, G. et al. TREM2 mutations implicated in neurodegeneration impair cell surface transport and phagocytosis. Sci. Transl. Med. 6, 243ra86 (2014).

20. Xiang, X. et al. TREM2 deficiency reduces the efficacy of immunotherapeutic amyloid clearance. EMBO Mol. Med. 8, 992–1004 (2016).

21. Mazaheri, F. et al. TREM2 deficiency impairs chemotaxis and microglial responses to neuronal injury. EMBO Rep. 18, 1186–1198 (2017).

22. Schlepckow, K., et al. Enhancing protective microglial activities with a dual function TREM2 antibody to the stalk region. EMBO Mol. Med. 12, e11227 (2020).

23. Price, B. R. et al. Therapeutic Trem2 activation ameliorates amyloid-beta deposition and improves cognition in the 5XFAD model of amyloid deposition. J. Neuroinflammation 17, 238 (2020).

24. van Lengerich, B. et al. A TREM2-activating antibody with a blood-brain barrier transport vehicle enhances microglial metabolism in Alzheimer’s disease models. Nat. Neurosci. 26, 416–429 (2023).

25. Wang, L. et al. Single-cell reconstruction of the adult human heart during heart failure and recovery reveals the cellular landscape underlying cardiac function. Nat. Cell Biol. 22, 108–119 (2020).

26. Udayar, V., Chen, Y., Sidransky, E. & Jagasia, R. Lysosomal dysfunction in neurodegeneration: emerging concepts and methods. Trends Neurosci. 45, 184–199 (2022).

27. Baker, M. et al. Mutations in progranulin cause tau-negative frontotemporal dementia linked to chromosome 17. Nature 442, 916–919 (2006).

28. Cruts, M. et al. Null mutations in progranulin cause ubiquitin-positive frontotemporal dementia linked to chromosome 17q21. Nature 442, 920–924 (2006).

29. Gass, J. et al. Mutations in progranulin are a major cause of ubiquitin-positive frontotemporal lobar degeneration. Hum. Mol. Genet. 15, 2988–3001 (2006).

30. Smith, K. R. et al. Strikingly different clinicopathological phenotypes determined by progranulin-mutation dosage. Am. J. Hum. Genet. 90, 1102– 1107 (2012).

31. Götzl, J. K. et al. Common pathobiochemical hallmarks of progranulin-associated frontotemporal lobar degeneration and neuronal ceroid lipofuscinosis. Acta Neuropathol. 127, 845–860 (2014).

32. Tanaka, Y. et al. Progranulin regulates lysosomal function and biogenesis through acidification of lysosomes. Hum. Mol. Genet. 26, 969–988 (2017).

33. Martens, L. H. et al. Progranulin deficiency promotes neuroinflammation and neuron loss following toxin-induced injury. J. Clin. Invest. 132, (2022).

34. Van Deerlin, V. M. et al. Common variants at 7p21 are associated with frontotemporal lobar degeneration with TDP-43 inclusions. Nat. Genet. 42, 234–239 (2010).

35. Cruchaga, C. et al. Association of TMEM106B gene polymorphism with age at onset in granulin mutation carriers and plasma granulin protein levels. Arch. Neurol. 68, 581–586 (2011).

36. Finch, N. et al. TMEM106B regulates progranulin levels and the penetrance of FTLD in GRN mutation carriers. Neurology 76, 467–474 (2011).

37. van der Zee, J. et al. TMEM106B is associated with frontotemporal lobar degeneration in a clinically diagnosed patient cohort. Brain 134, 808–815 (2011).

38. Feng, T. et al. Loss of TMEM106B and PGRN leads to severe lysosomal abnormalities and neurodegeneration in mice. EMBO Rep. 21, e50219 (2020).

39. Werner, G. et al. Loss of TMEM106B potentiates lysosomal and FTLD-like pathology in progranulin-deficient mice. EMBO Rep. 21, e50241 (2020).

40. Zhou, X. et al. Loss of Tmem106b exacerbates FTLD pathologies and causes motor deficits in progranulin-deficient mice. EMBO Rep. 21, e50197 (2020).

41. Hüttenrauch, M. et al. Glycoprotein NMB: a novel Alzheimer’s disease associated marker expressed in a subset of activated microglia. Acta Neuropathol. Commun. 6, 108 (2018).

42. Gazestani, V. et al. Early Alzheimer’s disease pathology in human cortex involves transient cell states. Cell 186, 4438–4453.e23 (2023).

43. Martins-Ferreira, R. et al. The Human Microglia Atlas (HuMicA) unravels changes in disease-associated microglia subsets across neurodegenerative conditions. Nat. Commun. 16, 739 (2025).

44. Wallings, R. L. et al. ASO-mediated knock-down of GPNMB in mutant-GRN and in Grn-deficient peripheral myeloid cells disrupts lysosomal function and immune responses. Mol. Neurodegener. 20, 41 (2025).

45. Caredio, D. et al. Gpnmb defines a phagocytic state of microglia linked to cell death in prion disease mouse model. Nat. Commun. 17, (2026).

46. Gillett, D. A., Wallings, R. L., Uriarte Huarte, O. & Tansey, M. G. Progranulin and GPNMB: interactions in endo-lysosome function and inflammation in neurodegenerative disease. J. Neuroinflammation 20, 286 (2023).

47. Tejwani, L. et al. Progranulin deficiency-induced lysosomal dysfunction drives maladaptive myeloid cell states through the MITF/TFE transcription factors. Immunity 59, 2164–2183.e11 (2026).

48. Balak, C. D. et al. Lysosomal dysfunction drives a transcriptional and epigenetic signature found in disease-associated microglia in neurodegenerative diseases. Immunity 59, 2142–2163.e8 (2026).

49. Lee, D. et al. Plasticity of human microglia and brain perivascular macrophages in aging and Alzheimer’s disease. Nat. Genet. 58, 2152–2163 (2026).

50. Saftig, P. et al. Mice deficient for the lysosomal proteinase cathepsin D exhibit progressive atrophy of the intestinal mucosa and profound destruction of lymphoid cells. EMBO J. 14, 3599–3608 (1995).

51. Sturchler-Pierrat, C. et al. Two amyloid precursor protein transgenic mouse models with Alzheimer disease-like pathology. Proc Natl Acad Sci USA 94, 13287–13292 (1997).

52. Radde, R. et al. Abeta42-driven cerebral amyloidosis in transgenic mice reveals early and robust pathology. EMBO Rep. 7, 940–946 (2006).

53. Colombo, A. et al. Loss of NPC1 enhances phagocytic uptake and impairs lipid trafficking in microglia. Nat. Commun. 12, 1158 (2021).

54. Reich, M. et al. Peripheral expression of brain-penetrant progranulin rescues pathologies in mouse models of frontotemporal lobar degeneration. Sci. Transl. Med. 16, eadj7308 (2024).

55. Bacioglu, M. et al. Neurofilament light chain in blood and CSF as marker of disease progression in mouse models and in neurodegenerative diseases. Neuron 91, 56–66 (2016).

56. Ashton, N. J. et al. A multicentre validation study of the diagnostic value of plasma neurofilament light. Nat. Commun. 12, 3400 (2021).

57. Xiang, X. et al. Microglial activation states drive glucose uptake and FDG-PET alterations in neurodegenerative diseases. Sci. Transl. Med. 13, eabe5640 (2021).

58. Zeisel, A. et al. Molecular architecture of the mouse nervous system. Cell 174, 999–1014.e22 (2018).

59. Ximerakis, M. et al. Single-cell transcriptomic profiling of the aging mouse brain. Nat. Neurosci. 22, 1696–1708 (2019).

60. Pesämaa, I. et al. A microglial activity state biomarker panel differentiates FTD-granulin and Alzheimer’s disease patients from controls. Mol. Neurodegener. 18, 70 (2023).

61. Bogacki, E. C. et al. GPNMB is a biomarker for lysosomal dysfunction and is secreted via LRRK2-modulated lysosomal exocytosis. Sci. Adv. 11, eadv1434 (2025).

62. De Schepper, S. et al. Perivascular cells induce microglial phagocytic states and synaptic engulfment via SPP1 in mouse models of Alzheimer’s disease. Nat. Neurosci. 26, 406–415 (2023).

63. Kana, V. et al. CSF-1 controls cerebellar microglia and is required for motor function and social interaction. J. Exp. Med. 216, 2265–2281 (2019).

64. Obst, J. et al. Inhibition of IL-34 Unveils Tissue-Selectivity and Is Sufficient to Reduce Microglial Proliferation in a Model of Chronic Neurodegeneration. Front. Immunol. 11, 579000 (2020).

65. Zhan, L. et al. A MAC2-positive progenitor-like microglial population is resistant to CSF1R inhibition in adult mouse brain. eLife 9, (2020).

66. Cantuti-Castelvetri, L. et al. Defective cholesterol clearance limits remyelination in the aged central nervous system. Science 359, 684–688 (2018).

67. Burns, J. C. et al. Differential accumulation of storage bodies with aging defines discrete subsets of microglia in the healthy brain. eLife 9, (2020).

68. Baumer, Y. et al. Ultramorphological analysis of plaque advancement and cholesterol crystal formation in Ldlr knockout mouse atherosclerosis. Atherosclerosis 287, 100–111 (2019).

69. Baumer, Y., Irei, J. & Boisvert, W. A. Cholesterol crystals in the pathogenesis of atherosclerosis. Nat. Rev. Cardiol. 22, 315–332 (2025).

70. Dinkel, L. et al. Myeloid cell-specific loss of NPC1 in mice recapitulates microgliosis and neurodegeneration in patients with Niemann-Pick type C disease. Sci. Transl. Med. 16, eadl4616 (2024).

71. Gabandé-Rodríguez, E. et al. Lipid-induced lysosomal damage after demyelination corrupts microglia protective function in lysosomal storage disorders. EMBO J. 38, (2019).

72. Wu, Y. et al. Microglial lysosome dysfunction contributes to white matter pathology and TDP-43 proteinopathy in GRN-associated FTD. Cell Rep. 36, 109581 (2021).

73. Depp, C. et al. Myelin dysfunction drives amyloid-β deposition in models of Alzheimer’s disease. Nature 618, 349–357 (2023).

74. Suzuki, C. et al. Lack of Cathepsin D in the central nervous system results in microglia and astrocyte activation and the accumulation of proteinopathy-related proteins. Sci. Rep. 12, 11662 (2022).

75. Roy, E. R. et al. Type I interferon response drives neuroinflammation and synapse loss in Alzheimer disease. J. Clin. Invest. 130, 1912–1930 (2020).

76. Wang, S. et al. TREM2 drives microglia response to amyloid-β via SYK-dependent and -independent pathways. Cell 185, 4153–4169.e19 (2022).

77. Mangalmurti, A. et al. Microglial CLEC7A restrains amyloid beta plaque pathology in a mouse model of Alzheimer’s disease. Alzheimers Dement 21, e70943 (2025).

78. Reifschneider, A. et al. Loss of TREM2 rescues hyperactivation of microglia, but not lysosomal deficits and neurotoxicity in models of progranulin deficiency. EMBO J. 41, e109108 (2022).

79. Lee, S.-H. et al. Trem2 restrains the enhancement of tau accumulation and neurodegeneration by β-amyloid pathology. Neuron 109, 1283–1301.e6 (2021).

80. Sogorb-Esteve, A. et al. Proteomic analysis reveals distinct cerebrospinal fluid signatures across genetic frontotemporal dementia subtypes. Sci. Transl. Med. 17, eadm9654 (2025).

81. Eninger, T. et al. Signatures of glial activity can be detected in the CSF proteome. Proc Natl Acad Sci USA 119, e2119804119 (2022).

82. Lazaratos, A.-M., Annis, M. G. & Siegel, P. M. GPNMB: a potent inducer of immunosuppression in cancer. Oncogene 41, 4573–4590 (2022).

83. Meng, Y. et al. GPNMB+ Gal-3+ hepatic parenchymal cells promote immunosuppression and hepatocellular carcinogenesis. EMBO J. 42, e114060 (2023).

84. Safaiyan, S. et al. White matter aging drives microglial diversity. Neuron 109, 1100–1117.e10 (2021).

85. Berghoff, S. A. et al. Microglia facilitate repair of demyelinated lesions via post-squalene sterol synthesis. Nat. Neurosci. 24, 47–60 (2021).

86. Lawrence, A. R. et al. Microglia maintain structural integrity during fetal brain morphogenesis. Cell 187, 962–980.e19 (2024).

87. Ayata, P. et al. Lymphoid gene expression supports neuroprotective microglia function. Nature 648, 157–165 (2025).

88. Fumagalli, L. et al. Microglia heterogeneity, modeling and cell-state annotation in development and neurodegeneration. Nat. Neurosci. 28, 1381–1392 (2025).

89. Dolan, M.-J. et al. Exposure of iPSC-derived human microglia to brain substrates enables the generation and manipulation of diverse transcriptional states in vitro. Nat. Immunol. 24, 1382–1390 (2023).

90. Dagher, N. N. et al. Colony-stimulating factor 1 receptor inhibition prevents microglial plaque association and improves cognition in 3xTg-AD mice. J. Neuroinflammation 12, 139 (2015).

91. Casali, B. T., MacPherson, K. P., Reed-Geaghan, E. G. & Landreth, G. E. Microglia depletion rapidly and reversibly alters amyloid pathology by modification of plaque compaction and morphologies. Neurobiol. Dis. 142, 104956 (2020).

92. Spangenberg, E. et al. Sustained microglial depletion with CSF1R inhibitor impairs parenchymal plaque development in an Alzheimer’s disease model. Nat. Commun. 10, 3758 (2019).

93. Prabata, A., Ikeda, K., Rahardini, E. P., Hirata, K.-I. & Emoto, N. GPNMB plays a protective role against obesity-related metabolic disorders by reducing macrophage inflammatory capacity. J. Biol. Chem. 297, 101232 (2021).

94. Nickl, B., Qadri, F. & Bader, M. Anti-inflammatory role of Gpnmb in adipose tissue of mice. Sci. Rep. 11, 19614 (2021).

95. Götzl, J. K. et al. Early lysosomal maturation deficits in microglia triggers enhanced lysosomal activity in other brain cells of progranulin knockout mice. Mol. Neurodegener. 13, 48 (2018).

96. Götzl, J. K. et al. Opposite microglial activation stages upon loss of PGRN or TREM2 result in reduced cerebral glucose metabolism. EMBO Mol. Med. 11, (2019).

97. Huang, M. et al. Network analysis of the progranulin-deficient mouse brain proteome reveals pathogenic mechanisms shared in human frontotemporal dementia caused by GRN mutations. Acta Neuropathol. Commun. 8, 163 (2020).

98. Gillett, D. A., Neighbarger, N. K., Cole, C., Wallings, R. L. & Tansey, M. G. Investigating the role and regulation of GPNMB in progranulin-deficient macrophages. Front. Immunol. 15, 1417836 (2024).

99. Suda, M. et al. Glycoprotein nonmetastatic melanoma protein B regulates lysosomal integrity and lifespan of senescent cells. Sci. Rep. 12, 6522 (2022).

100. Liu, M. et al. GPNMB and ATP6V1A interact to mediate microglia phagocytosis of multiple types of pathological particles. Cell Rep. 44, 115343 (2025).

101. Johnson, E. C. B. et al. Large-scale proteomic analysis of Alzheimer’s disease brain and cerebrospinal fluid reveals early changes in energy metabolism associated with microglia and astrocyte activation. Nat. Med. 26, 769–780 (2020).

102. Pereira, J. B. et al. Microglial activation protects against accumulation of tau aggregates in nondemented individuals with underlying Alzheimer’s disease pathology. *Nat*. Aging 2, 1138–1144 (2022).

103. Qiu, Y. et al. Definition of the contribution of an Osteopontin-producing CD11c+ microglial subset to Alzheimer’s disease. Proc Natl Acad Sci USA 120, e2218915120 (2023).

104. Johnson, E. C. B. et al. Cerebrospinal fluid proteomics define the natural history of autosomal dominant Alzheimer’s disease. Nat. Med. 29, 1979–1988 (2023).

105. Mancuso, R. et al. Xenografted human microglia display diverse transcriptomic states in response to Alzheimer’s disease-related amyloid-β pathology. Nat. Neurosci. 27, 886–900 (2024).

106. Zigdon, H. et al. Identification of a biomarker in cerebrospinal fluid for neuronopathic forms of Gaucher disease. PLoS ONE 10, e0120194 (2015).

107. Kramer, G. et al. Elevation of glycoprotein nonmetastatic melanoma protein B in type 1 Gaucher disease patients and mouse models. FEBS Open Bio 6, 902–913 (2016).

108. Murugesan, V. et al. Validating glycoprotein non-metastatic melanoma B (gpNMB, osteoactivin), a new biomarker of Gaucher disease. Blood Cells Mol. Dis. 68, 47–53 (2018).

109. Moloney, E. B., Moskites, A., Ferrari, E. J., Isacson, O. & Hallett, P. J. The glycoprotein GPNMB is selectively elevated in the substantia nigra of Parkinson’s disease patients and increases after lysosomal stress. Neurobiol. Dis. 120, 1–11 (2018).

110. Srinivasan, K. et al. Untangling the brain’s neuroinflammatory and neurodegenerative transcriptional responses. Nat. Commun. 7, 11295 (2016).

111. Zhu, S. et al. Targeted multiple reaction monitoring analysis of CSF identifies UCHL1 and GPNMB as candidate biomarkers for ALS. J. Mol. Neurosci. 69, 643–657 (2019).

112. Brendel, M. et al. Glial Activation and Glucose Metabolism in a Transgenic Amyloid Mouse Model: A Triple-Tracer PET Study. J. Nucl. Med. 57, 954–960 (2016).

113. Sobue, A. et al. Microglial gene signature reveals loss of homeostatic microglia associated with neurodegeneration of Alzheimer’s disease. Acta Neuropathol. Commun. 9, 1 (2021).

## Method references

1. Spangenberg, E. E. et al. Eliminating microglia in Alzheimer’s mice prevents neuronal loss without modulating amyloid-β pathology. Brain 139, 1265–1281 (2016).

2. Brendel, M. et al. Glial Activation and Glucose Metabolism in a Transgenic Amyloid Mouse Model: A Triple-Tracer PET Study. J. Nucl. Med. 57, 954–960 (2016).

3. Gnörich, J. et al. Depletion and activation of microglia impact metabolic connectivity of the mouse brain. J. Neuroinflammation 20, 47 (2023).

4. Zeisel, A. et al. Molecular architecture of the mouse nervous system. Cell 174, 999–1014.e22 (2018).

5. Ximerakis, M. et al. Single-cell transcriptomic profiling of the aging mouse brain. Nat. Neurosci. 22, 1696–1708 (2019).

6. Liberzon, A. et al. The Molecular Signatures Database (MSigDB) hallmark gene set collection. Cell Syst. 1, 417–425 (2015).

7. Keren-Shaul, H. et al. A Unique Microglia Type Associated with Restricting Development of Alzheimer’s Disease. Cell 169, 1276–1290.e17 (2017).

8. Silvin, A. et al. Dual ontogeny of disease-associated microglia and disease inflammatory macrophages in aging and neurodegeneration. Immunity 55, 1448–1465.e6 (2022).

9. Sobue, A. et al. Microglial gene signature reveals loss of homeostatic microglia associated with neurodegeneration of Alzheimer’s disease. Acta Neuropathol. Commun. 9, 1 (2021).

10. Fumagalli, L. et al. Microglia heterogeneity, modeling and cell-state annotation in development and neurodegeneration. Nat. Neurosci. 28, 1381–1392 (2025).

11. Depp, C. et al. Myelin dysfunction drives amyloid-β deposition in models of Alzheimer’s disease. Nature 618, 349–357 (2023).

12. van Lengerich, B. et al. A TREM2-activating antibody with a blood-brain barrier transport vehicle enhances microglial metabolism in Alzheimer’s disease models. Nat. Neurosci. 26, 416–429 (2023).

13. Lee, S.-H. et al. Trem2 restrains the enhancement of tau accumulation and neurodegeneration by β-amyloid pathology. Neuron 109, 1283–1301.e6 (2021).

14. Reich, M. et al. Peripheral expression of brain-penetrant progranulin rescues pathologies in mouse models of frontotemporal lobar degeneration. Sci. Transl. Med. 16, eadj7308 (2024).

15. Nott, A. et al. Brain cell type-specific enhancer-promoter interactome maps and disease-risk association. Science 366, 1134–1139 (2019).

16. Nott, A., Schlachetzki, J. C. M., Fixsen, B. R. & Glass, C. K. Nuclei isolation of multiple brain cell types for omics interrogation. Nat. Protoc. 16, 1629–1646 (2021).

17. Heinz, S. et al. Simple combinations of lineage-determining transcription factors prime cis-regulatory elements required for macrophage and B cell identities. Mol. Cell 38, 576–589 (2010).

18. Texari, L. et al. An optimized protocol for rapid, sensitive and robust on-bead ChIP-seq from primary cells. STAR Protocols 2, 100358 (2021).

19. Langmead, B. & Salzberg, S. L. Fast gapped-read alignment with Bowtie 2. Nat. Methods 9, 357–359 (2012).

20. Fixsen, B. R. et al. SALL1 enforces microglia-specific DNA binding and function of SMADs to establish microglia identity. Nat. Immunol. 24, 1188–1199 (2023).

21. Kislinger, G. et al. Neurons on tape: Automated Tape Collecting Ultramicrotomy-mediated volume EM for targeting neuropathology. Methods Cell Biol. 177, 125– 170 (2023).

22. Schindelin, J. et al. Fiji: an open-source platform for biological-image analysis. Nat. Methods 9, 676–682 (2012).

23. Lourbopoulos, A. et al. An improved method for sampling and quantitative protein analytics of cerebrospinal fluid of individual mice. Mol. Cell. Proteomics 24, 100958 (2025).

24. Demichev, V. et al. dia-PASEF data analysis using FragPipe and DIA-NN for deep proteomics of low sample amounts. Nat. Commun. 13, 3944 (2022).

25. Wefers, B., Wurst, W. & Kühn, R. Gene editing in mouse zygotes using the crispr/cas9 system. Methods Mol. Biol. 2631, 207–230 (2023).

26. Concordet, J.-P. & Haeussler, M. CRISPOR: intuitive guide selection for CRISPR/Cas9 genome editing experiments and screens. Nucleic Acids Res. 46, W242–W245 (2018).

27. Dobin, A. et al. STAR: ultrafast universal RNA-seq aligner. Bioinformatics 29, 15– 21 (2013).

28. Martin, M. Cutadapt removes adapter sequences from high-throughput sequencing reads. EMBnet j. 17, 10 (2011).

29. Hao, Y. et al. Integrated analysis of multimodal single-cell data. Cell 184, 3573–3587. (2021).

30. McGinnis, C. S., Murrow, L. M. & Gartner, Z. J. DoubletFinder: Doublet Detection in Single-Cell RNA Sequencing Data Using Artificial Nearest Neighbors. Cell Syst. 8, 329–337.e4 (2019).

31. Korsunsky, I. et al. Fast, sensitive and accurate integration of single-cell data with Harmony. Nat. Methods 16, 1289–1296 (2019).

32. Love, M. I., Huber, W. & Anders, S. Moderated estimation of fold change and dispersion for RNA-seq data with DESeq2. Genome Biol. 15, 550 (2014).

33. Hänzelmann, S., Castelo, R. & Guinney, J. GSVA: gene set variation analysis for microarray and RNA-seq data. BMC Bioinformatics 14, 7 (2013).

34. Yu, G., Wang, L.-G., Han, Y. & He, Q.-Y. clusterProfiler: an R package for comparing biological themes among gene clusters. OMICS 16, 284–287 (2012).

35. Aibar, S. et al. SCENIC: single-cell regulatory network inference and clustering. Nat. Methods 14, 1083–1086 (2017).

36. Aboyoun, P., et al. NanoStringNCTools: NanoString nCounter Tools. R Package Version 1.0.0. (2021).

